# Assembly mechanism and cryoEM structure of RecA recombination nucleofilaments from *Streptococcus pneumoniae*

**DOI:** 10.1101/2022.07.28.501814

**Authors:** Hertzog Maud, Perry Thomas Noé, Dupaigne Pauline, Serres Sandra, Morales Violette, Soulet Anne-Lise, C Bell Jason, Margeat Emmanuel, Kowalczykowski Stephen, Eric Le Cam, Fronzes Rémi, Polard Patrice

## Abstract

RecA-mediated Homologous Recombination (HR) is a key mechanism for genome maintenance and plasticity in bacteria. It proceeds through RecA assembly into a dynamic filament on ssDNA, the presynaptic filament, which mediates DNA homology search and ordered DNA strand exchange. Here, we combined structural, single molecule and biochemical approaches to characterize the ATP-dependent assembly mechanism of the presynaptic filament of RecA from *Streptococcus pneumoniae* (*Sp*RecA), in comparison to the *Escherichia coli* RecA (*Ec*RecA) paradigm. *Ec*RecA polymerization on ssDNA is assisted by the Single-Stranded DNA Binding (SSB) protein, which unwinds ssDNA secondary structures that block *Ec*RecA nucleofilament growth. We report that neither of the two paralogous pneumococcal SSBs could assist *Sp*RecA polymerization on ssDNA. Instead, we found that the conserved RadA helicase promotes this *Sp*RecA nucleofilamentation in an ATP-dependent manner. This allowed us to solve the atomic structure of such a long native *Sp*RecA nucleopolymer by cryoEM stabilized with ATPγS. It was found to be equivalent to the crystal structure of the *Ec*RecA filament with a marked difference in how RecA mediates nucleotide orientation in the stretched ssDNA. Then, our results show that *Sp*RecA and *Ec*RecA HR activities are different, in correlation with their distinct ATP-dependent ssDNA binding modes.

## Introduction

Homologous recombination (HR) is a DNA strands exchange process essential for multiple pathways of genome maintenance and plasticity in all kingdoms of life (Cox, 2007; Kowalczykowski, 2016). Defects in any of these pathways lead to deleterious consequences, such as cell death or various types of cancer (Liu et al., 2011). HR relies on the pairing of a single-stranded DNA (ssDNA) molecule with one complementary strand in a double-stranded DNA (dsDNA) to generate a three-stranded DNA structure, commonly referred to as a synaptic product or a D-loop structure (for <u>D</u>isplacement-<u>loop</u>) (Michel and Leach, 2012). This reaction is catalyzed by a widespread and conserved group of enzymes, defined hereafter as HR recombinases and named RecA in bacteria, Rad51/Dmc1 in eukaryotes, RadA in archaea. They form the RecA/Rad51 protein family, unified by a conserved ATP binding and hydrolysis core domain and a common HR mechanism (Bell and Kowalczykowski, 2016). They promote the pairing and exchange of homologous DNA molecules by polymerizing first on a ssDNA molecule to generate the so-called presynaptic nucleofilament (Kowalczykowski, 2015). Once assembled, the presynaptic nucleofilament scans for an homologous sequence in dsDNA and promotes ssDNA pairing with a complementary DNA sequence to generate the D-loop (Renkawitz et al., 2014). The assembly and disassembly of RecA/Rad51 nucleofilaments is finely tuned by the binding and hydrolysis of nucleotides at the interface between monomers in the polymer. As such, RecA/Rad51 nucleofilaments are dynamic and the regulation of the DNA-dependent NTP binding and hydrolysis cycle is at the heart of the HR process.

RecA from *Escherichia coli* (*Ec*) is a model protein for the bacterial HR recombinases (Del Val, 2019). Crystal structures of a chimera made of 6 or 5 fused protomers of *Ec*RecA truncated for N-terminal (1-30) and C-terminal (336–353) residues and bound to DNA in the presence of a non-hydrolyzable ATP derivative have revealed many key features about the organization of presynaptic filament assembly and the mechanism of its pairing with a complementary ssDNA molecule (Chen et al., 2008). First, ssDNA and ATP bind to RecA-RecA interfaces cooperatively, explaining the ATP dependency for RecA polymerization on ssDNA. Second, the γ-phosphate of ATP is sensed across the RecA-RecA interface by two lysine residues that stimulate ATP hydrolysis, providing a mechanism for DNA release. Third, the nucleoprotein filament adopts a right-handed helical shape with six *Ec*RecA monomers per turn, which stretch the ssDNA about 1.6-fold in comparison with a B-form dsDNA. Remarkably, the ssDNA is organized in the filament in regularly separated B-form triplets of nucleotides, including two bases exposed externally that restricts the homology search to Watson–Crick-type base pairing within a recipient DNA. In addition, using a chimera of 9 fused *Ec*RecA subunits, Yang et al. recently solved by high-resolution cryoEM the structure of a D-loop assembled by this chimeric *Ec*RecA polymer in the presence of a non-hydrolysable ATP derivative. This has revealed a small loop in the C-terminal region of *Ec*RecA, conserved in other bacterial RecA, which helps to open the recipient dsDNA. Also, the displaced non-complementary strand of the recipient dsDNA in the nucleoprotein D-loop structure is bound by a secondary DNA-binding site in *Ec*RecA (Yang et al., 2020). In parallel to these advances in the structural organization of *Ec*RecA nucleofilaments HR intermediates, the development of single molecule (SM) studies using fluorescently labelled *Ec*RecA revealed important aspects in the dynamics of its ATP-dependent polymerization on DNA (Bell and Kowalczykowski, 2016a). First, these studies highlighted the bidirectional growth of *Ec*RecA filaments along ssDNA, with a more rapid extension from 5’ to 3’ direction (Bell et al., 2012). Secondly, they provided a direct imaging of the slow initial *Ec*RecA interaction on ssDNA prior to the rapid growth of the nucleofilament, referred to as the nucleation and extension stages, respectively (Joo et al., 2006). Third, these SM studies unveiled key features of the DNA homology search and subsequent pairing stages mediated by *Ec*RecA presynaptic filament, which proceeds through an inchworm mechanism and a 3-nucleotide stepping during DNA strand exchange, respectively (Chen et al., 2008). Similar structural and SM analysis conducted on Rad51/DMC1 pointed at the conservation of *Ec*RecA properties in eukaryotic HR recombinases, pointing at their general character of these properties to all members of the RecA/Rad51 family (Lee et al., 2015).

Another common feature of HR recombinases is the assistance of accessory factors that modulate their polymerization/depolymerization from DNA templates and/or their DNA strands exchange activities (Antony and Lohman, 2019). These HR modulators are differentially conserved, with some being found in a whole kingdom of life and others limited to few species. They compose distinct and partially overlapping subsets of HR effectors that define specific pathways of genome maintenance and plasticity. In bacteria, a widely conserved HR effector is the SSB protein (<u>S</u>ingle-<u>S</u>tranded DNA <u>B</u>inding). SSB is firstly known as being essential to cell growth *via* its action at the replication forks where it protects ssDNA and assists its replication. Reconstitution of *E. coli* HR *in vitro* has highlighted three distinct roles of its cognate SSB in counteracting or assisting its DNA interacting activities (Bianco, 2017): one is to prevent RecA nucleation if bound first on the ssDNA; a second is to promote RecA polymerization along ssDNA, by removing the secondary structures that impede filament growth; a third is to bind to the extruded parental strand during the DNA strand exchange reaction, stabilizing the recombination product and favoring the incorporation of ssDNA (a step referred to as a DNA branch migration). Another key and conserved bacterial effector acting in these postsynaptic stages of HR is the RadA helicase, which has been found to facilitate ssDNA recombination from D-loop stuctures *via* interaction with RecA and by driving DNA branch migration (Cooper and Lovett, 2016) (Torres et al., 2019)(Marie et al. 2017).7/28/2022 4:39:00 AM Other known HR effectors act at one or more of these steps of the HR recombinase activity cycle through static or dynamic protein-protein interactions and/or DNA remodeling activities (Liu et al., 2011).

Not all bacterial RecAs exhibit the same intrinsic activities as *Ec*RecA. Notable examples are RecA from *Deinococcus radiodurans* (*Dr*) and from *Pseudomonas aeruginosa* (*Pa*). These two bacterial species undergo high HR rate in their natural environment to sustain their growth under extreme radiative or oxidative conditions, respectively (Baitin et al., 2006; Cox and Battista, 2005). *Dr*RecA is less homologous to *Ec*RecA than *Pa*RecA, i.e. 61% and 71% of sequence identity, respectively. Their DNA binding affinity was found higher than that of *Ec*RecA, along with a stronger ability to displace its cognate SSB from ssDNA for *Pa*RecA and with a reverse recombination from dsDNA to ssDNA for *Dr*RecA. Another reported deviation to the *Ec*RecA paradigm is RecA from *Streptococcus pneumoniae* (*Sp*, the pneumococcus), which is well known to undergo high HR rate during the process of genetic transformation induced in response to multiple stress during the state of competence (Johnston et al., 2014). Previous biochemical studies comparing *Sp*RecA and *Ec*RecA activities highlighted two marked differences between those two HR recombinases, which share 63% of identity. First, *Sp*RecA was found intrinsically less efficient than *Ec*RecA in directing DNA strand exchange between a long circular ssDNA with a complementary strand in a linear duplex DNA; second, *Sp*RecA interaction with ssDNA appears to be negatively challenged by any of its two cognate paralogous SSB proteins, namely SsbA and SsbB, the former being essential and involved in DNA replication and genome maintenance processes and the latter being restrictively expressed during competence and involved in the mechanism of natural transformation (Attaiech et al., 2011; Grove and Bryant, 2006). What precisely determine these functional deviations of *Sp*RecA in comparison with *Ec*RecA remains enigmatic.

Here, we combined classical biochemical techniques with single-molecule and structural approaches to closely examine the DNA interacting properties of *Sp*RecA in comparison with *Ec*RecA. Together, these experiments highlight significant variations in the ATP-dependent dynamics and structure of the *Sp*RecA presynaptic filament, including the lack of assistance in its elongation by any SSB protein. Unexpectedly, however, we found that the *Sp*RadA helicase could promote such an extension of the *Sp*RecA presynaptic filament in an ATP-dependent manner. Altogether, this detailed analysis provides important molecular insights into the distinct efficiency of *Sp*RecA in catalyzing HR in comparison with *Ec*RecA.

## Results

### *Sp*RecA forms short presynaptic filaments

First, we purified *Sp*RecA and analyzed by transmission electron microscopy (TEM) it ATP-dependent polymerizing activity on ssDNA in comparison with purified *Ec*RecA, by using circular form of ΦX174 bacteriophage (5386 nucleotides long) as a template. *Sp*RecA added in saturating concentration to fully cover all ssDNA molecules forms dense structures on ssDNA in the presence of ATP (Figure1b). These structures result from *Sp*RecA binding to ssDNA as they are not observed with naked ssDNA (Figure 1a). The same experiment performed with an ATP regenerating system did not change the result (Figure 1c). By contrast, short polymers could be detected in the presence of the poorly hydrolysable ATPγS derivative (Figure 1d). We also observed similar extended filaments in the presence of ATP and of BeF_3_, a Pi analogue known to notably inhibit ATP hydrolysis (Figure 1e and Supplemental Figure 1). The same TEM analysis performed with *Ec*RecA showed that it also forms small dense structures on ssDNA (Figure 1f) in the presence of ATP. By contrast, *Ec*RecA was able to form extended filaments in the presence of ATP together with an ATP regenerating system (Figure 1g), showing not only its binding but also its assembly along ssDNA. Furthermore, *Ec*RecA appeared to polymerize extensively along ΦX174 ssDNA in the presence of ATPγS (Figure 1h), and for a 6-fold longer distance than *Sp*RecA in the same conditions (mean value of 767 nm and 122 nm, respectively; see Figure 1i). Both recombinases could generate several filaments on the same ΦX174 ssDNA molecule in experiments performed with ATPγS, indicative of several nucleation events for both recombinases followed by their polymerization. The total length of *Sp*RecA nucleofilaments per individual ssDNA molecule was ∼0.34 µm in the presence of ATPγS (and ∼0.47 µm in the presence of ATP and BeF_3_), contrasting with the ∼2.06 µm measured for *Ec*RecA in the presence of ATPγS. Also, in these conditions, *Ec*RecA does not fully cover the circular ssDNA template, indicating that its polymerization is blocked at some sites, most probably secondary structures that formed on ssDNA (Bell et al., 2012). Interestingly, *Sp*RecA seems to be less efficient than *Ec*RecA to unfold such structures, explaining why *Sp*RecA generate more and shorter filaments on the long ssDNA template. Altogether, these results revealed a marked difference between *Sp*RecA and *Ec*RecA in their ability to extend their nucleofilamentation under these stabilizing conditions that block ATP hydrolysis and their release from ssDNA.

**Figure 1.**
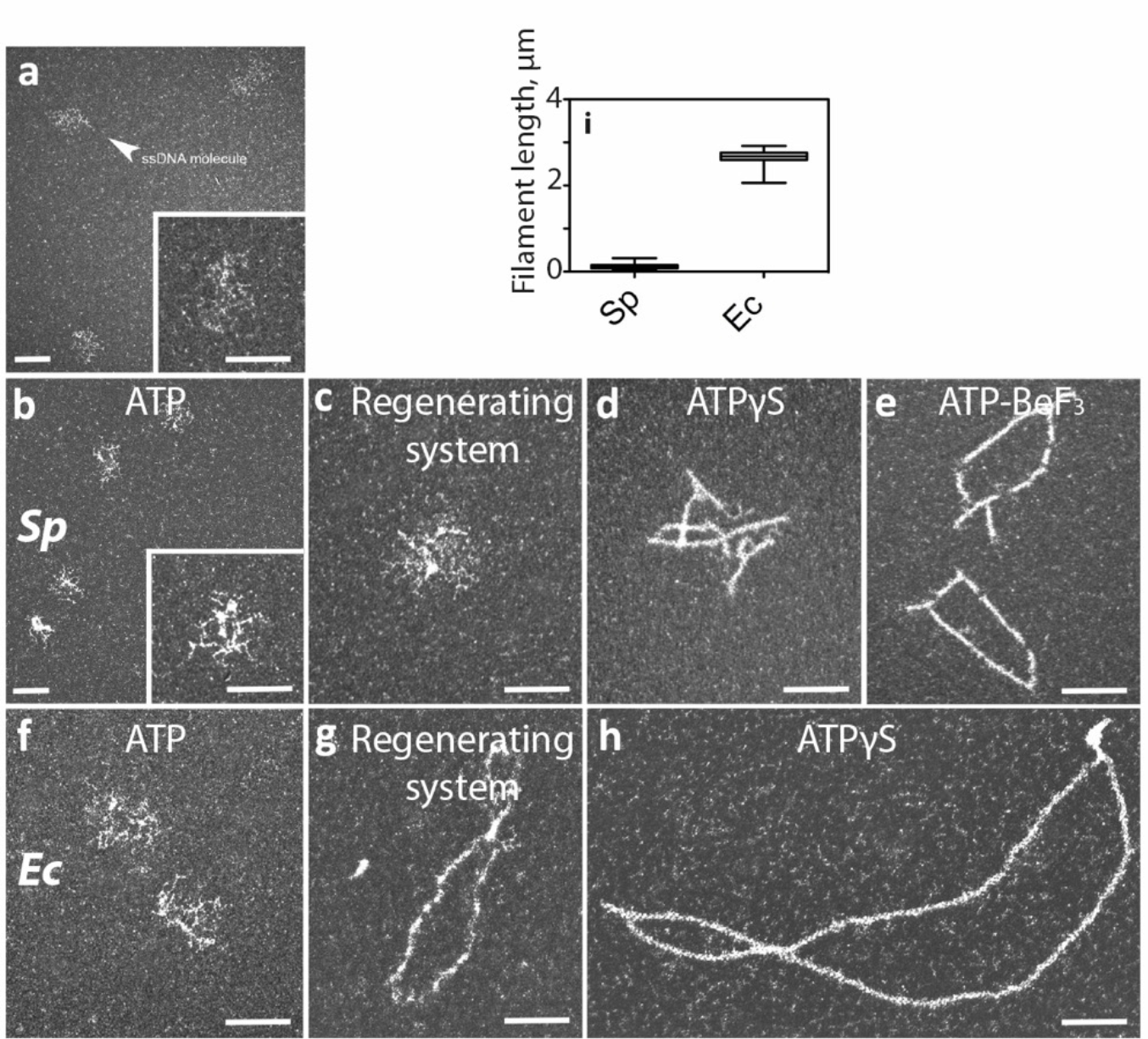
TEM analysis of *Sp*RecAand *Ec*RecA polymerisation on ssDNA. From a to h, representative electron micrographs images of ΦX ssDNA alone (a) or with *Sp*RecA and ATP (b), *Sp*RecA, ATP and an ATP regenerating system (c), *Sp*RecA and ATPgS (d), *Sp*RecA and ATP-BeF_3_ (e), with *Ec*RecA and ATP (f), RecA_*Ec*_, ATP and an ATP regenerating system (g), RecA_*Ec*_ and ATPgS (h). In i: measured length of the filaments made with *Ec*RecA or *Sp*RecA in presence of ATPgS. All scale bars represent 200 nm.

### *Sp*RecA polymerisation on ssDNA is not assisted by any SSB protein

*Ec*RecA presynaptic filamentation is assisted by its cognate SSB *via* its melting activity of secondary structures that form on ssDNA (Kowalczykowski et al., 1987). This has been generalized to all bacterial RecA similarly studied *in vitro* (Bianco, 2017). In *S. pneumoniae*, SSB appears to counteract the presynaptic HR step of *Sp*RecA, as indirectly evaluated by measuring the rate of ssDNA induced *Sp*RecA ATP hydrolysis, while still stimulating the subsequent DNA strand exchange step (Grove and Bryant, 2006). This negative competitive effect of SSB on *Sp*RecA interaction with ssDNA was observed with any of the two paralogous pneumococcal SSB, SsbA and SsbB, as well as with *Ec*SSB (Attaiech et al., 2011; Grove and Bryant, 2006; Nayak and Bryant, 2015; Steffen and Bryant, 2001). Conversely, SsbA and SsbB behave similarly as *Ec*SSB in stimulating *Ec*RecA ATPase and HR activities. These earlier studies pointed at a distinct HR activity of *Sp*RecA in comparison with *Ec*RecA, which is differently challenged by SSB proteins. We studied by TEM this interplay between *Sp*RecA and SSB proteins. Purified SSB proteins were used at a concentration allowing a full coverage of all ssDNA molecules. Typical images obtained by TEM at these saturating amounts of SsbA and SsbB are presented in Figure 2a and 2b, respectively, resulting in an identical pattern of interaction with the circular □X174 ssDNA template used. Subsequently, addition of either of these two SSB proteins to *Sp*RecA pre-bound to ssDNA in the presence of ATP alone or with an ATP regenerating system did not promote its polymerization but led to the same nucleocomplexes observed with the SSBs alone. In the same vein, addition of SsbA or SsbB to the short *Sp*RecA filaments formed in the presence of ATPγS did not promote their elongation but led to the binding of either of these two SSBs to the ssDNA unoccupied by *Sp*RecA (Figures 2c and 2e, respectively). By contrast, *Ec*SSB added to *Ec*RecA pre-incubated with ssDNA in the presence of ATPγS promotes a full coverage of the circular ssDNA molecules by *Ec*RecA (Figure 2d). The same result has been obtained by adding *Ec*SSB to *Ec*RecA incubated with ssDNA in the presence of ATP and an ATP regenerating system. Furthermore, we also found that *Ec*RecA polymerization along ssDNA could also be assisted by the two pneumococcal SSBs and, conversely, that *Ec*SSB failed to assist *Sp*RecA nucleofilamentation in any conditions. Altogether, these findings highlight that *Sp*RecA ATP-dependent filamentation on ssDNA is not assisted by any SSB, highlighting a main functional diversity? /deviation/disparity between *Sp*RecA and the *Ec*RecA paradigm and all other RecA proteins similarly studied so far.

**Figure 2.**
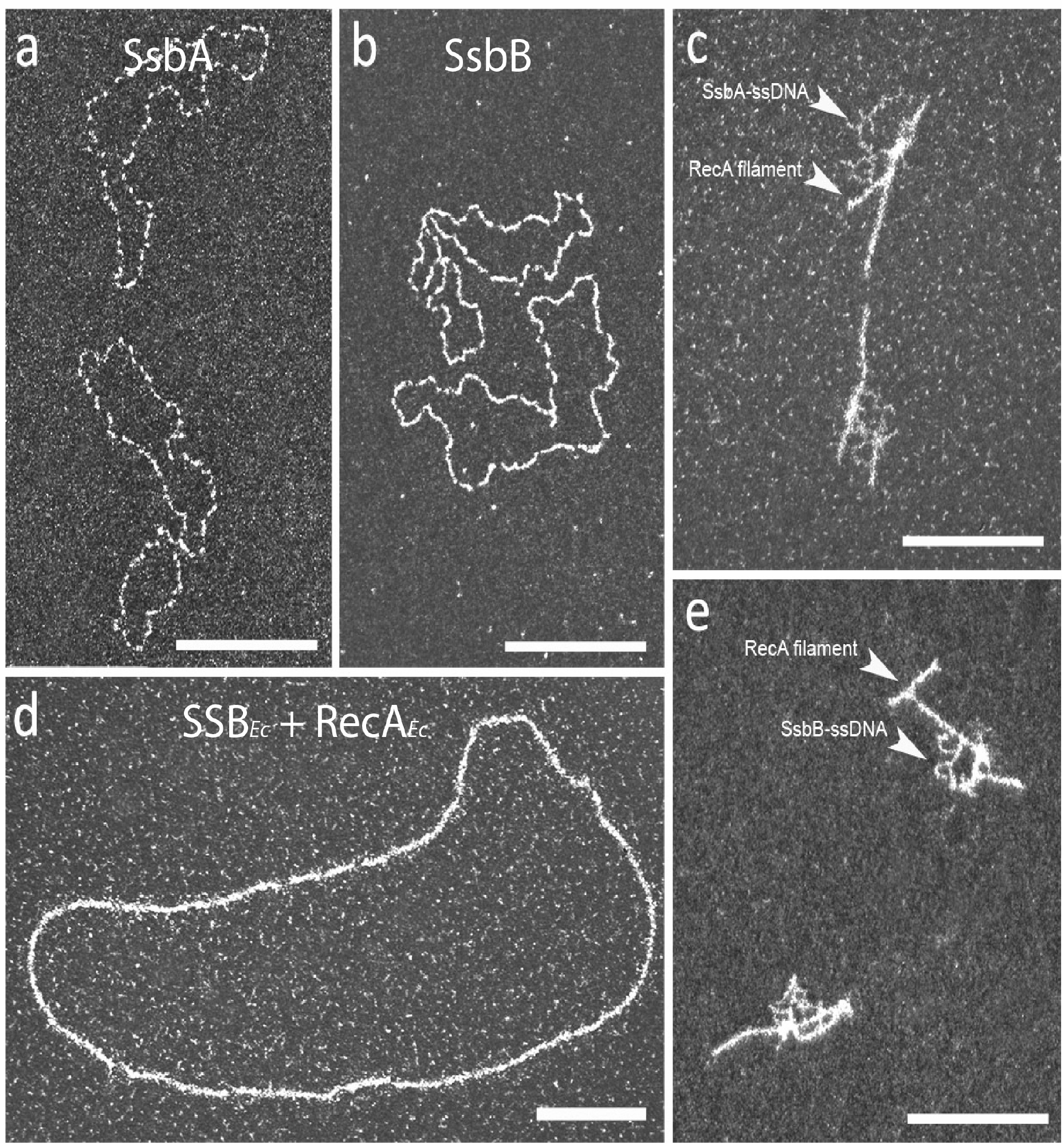
TEM analysis of *Sp*RecA and *Ec*RecA assembly on ssDNA in presence of *Sp*SsbA and *Sp*SsbB, or *Ec*SSB. Representative electron micrographs images of ΦX ssDNA incubated either with saturating amount of SsbA (a) or SsbB (b), or pre-incubated with *Sp*RecA and, next, incubated with saturating amount of SsbA (c) or SsbB (e), or pre-incubated with *Ec*RecA and, next, incubated with saturating amount of SSB (d). All scale bars represent 200 nm.

### The *Sp*RadA HR helicase extends *Sp*RecA polymerization on ssDNA

We next wondered whether *Sp*RecA nucleofilamentation on ssDNA could be assisted by another HR effector. An obvious candidate was the *Sp*RadA protein, which we found to interact with RecA and to be a DnaB-type hexameric helicase that canonically translocates along ssDNA fueled by ATP hydrolysis in the 5’ to 3’ direction (Marie et al., 2017). We investigated by TEM analysis whether *Sp*RadA could modulate *Sp*RecA extension along ssDNA, by using the circular form of the M13 bacteriophage. While a few small polymers were formed in the presence of hydrolysable ATP in the absence of *Sp*RadA, incubation of *Sp*RecA with *Sp*RadA in the presence of ATPγS promoted the formation of nucleofilaments 10-fold longer than those observed with *Sp*RecA alone in those conditions (Figure 3a and 3c). Sub-stoichiometric amounts of *Sp*RadA with respect to *Sp*RecA concentration (1:4) were sufficient to generate these polymers. Observation by negative staining showed that these longer filaments formed by mixing *Sp*RecA with *Sp*RadA are comparable to the helical filaments generated by *Sp*RecA alone, indicating that *Sp*RadA promoted *Sp*RecA polymerization extension along ssDNA. Next, we reproduced these experiments with *Sp*RadA^K101A^ point mutant, which was previously shown to be unable to hydrolyze ATP (Marie et al., 2017). This *Sp*RadA^K101A^ mutant, which still assembled into hexamers as wild-type protein (as visible on the EM grid; Figure 3b), was no longer able to promote the formation of long filaments when mixed with *Sp*RecA. This result shows that ATPγS hydrolysis is necessary to extend *Sp*RecA filamentation along ssDNA. The need for *Sp*RadA ATPase activity supports that it acts by translocating on the ssDNA to unwind the secondary structures that impede *Sp*RecA filament growth, but without physically blocking *Sp*RecA assembly on ssDNA as SSB does. However, we could not exclude that *Sp*RadA assists *Sp*RecA filamentation by another ATP-dependent mechanism relying on their interaction. Finally, based on this TEM analysis, we could not conclude whether *Sp*RadA is associated to these nucleofilaments.

**Figure 3.**
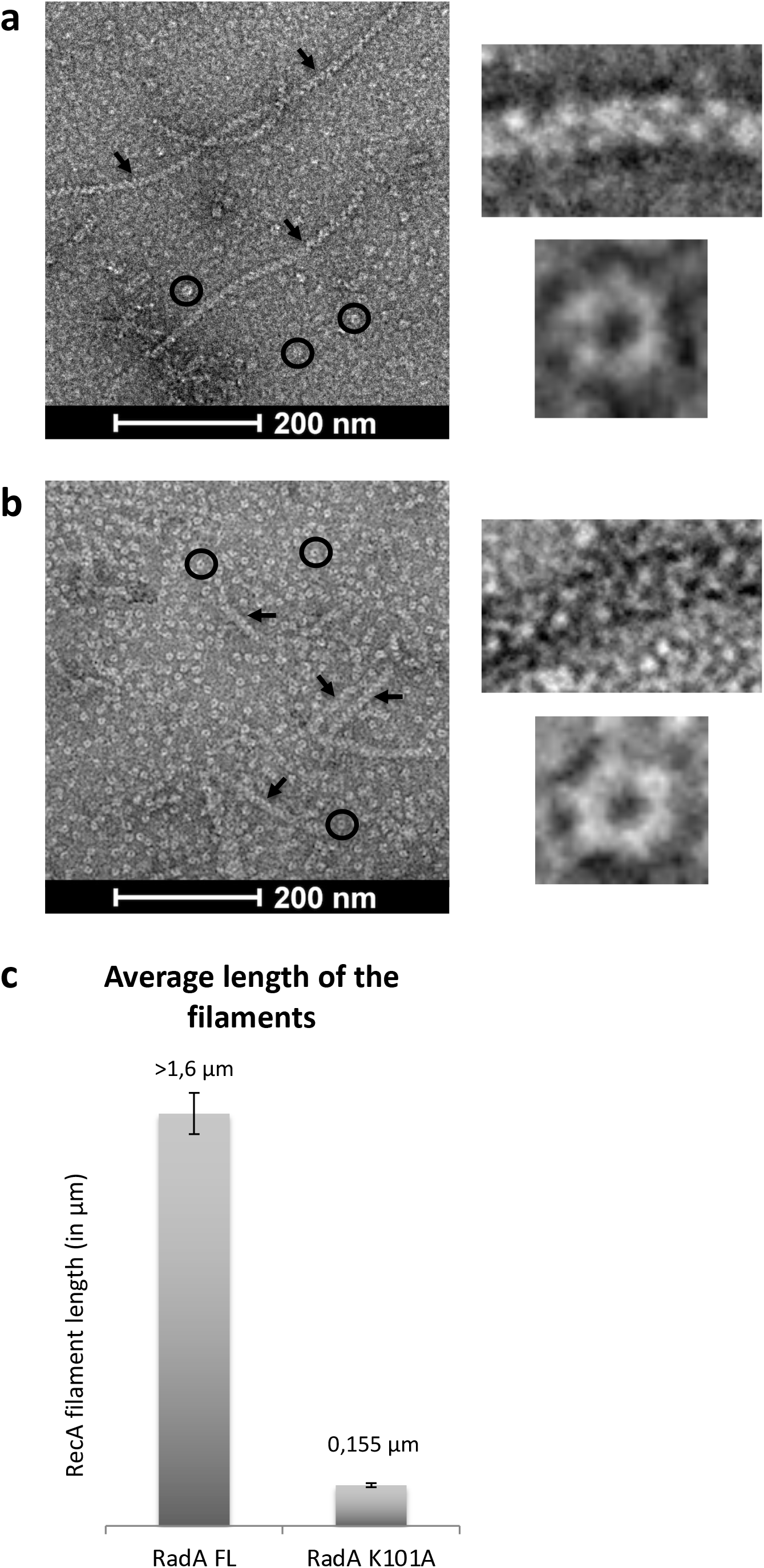
*Sp*RecA extends *Sp*RecA polymerization along ssDNA. a and b. Negatively stained EM images of presynaptic filaments in presence of RadA_FL_ (a) and of RadA_K101A_ (b), the presynaptic filaments and RadA proteins are shown by black arrows and circles respectively. The insets on the right show a zoom of presynaptic filaments and RadA_FL_ (a), and RadA_K101A_ (b) self-assembled into ring-shaped hexamers. c. Histogram of *Sp*RecA presynaptic filaments average length in the presence of RadA_FL_ and RadA_K101A_.

### CryoEM analysis of *Sp*RecA filaments assembled on ssDNA and dsDNA

*Sp*RecA nucleofilaments formed on long ssDNA molecules in the presence of ATPγS are too short to allow their structural analysis by cryoEM. This drawback has been overcome via the action of *Sp*RadA (see above). Long nucleoprotein filaments formed on M13 ssDNA were deposited on cryoEM Lacey grids and visualized using a 200 KeV Talos Arctica cryo-electron microscope (See Supplemental Figure 2a). We used helical reconstruction in Relion to obtain a 3.9Å resolution map. Using a non-symmetrized 3D reconstruction, we could determine in real space that these filaments display a helical symmetry and estimate the helical parameters (See Supplemental figure 3). These helical parameters were then imposed and refined during reconstruction to obtain the final map with 15.38 Å rise and 58.46 degrees of twist (corresponding to 6.16 subunits per turn of helix) (See methods and table 1). The final cryoEM map displayed key structural features of *Sp*RecA with the bulky side chains clearly visible (Supplemental Figure 2g). A homology structural model of the *Sp*RecA was obtained using Swissmodel using *Ec*RecA crystal structure as template. *Sp*RecA and *Ec*RecA proteins share 63% identity (sequence-based alignment) and we postulated that their structure should be similar in term of secondary structure elements and overall fold. This initial model was docked into the map and the *Sp*RecA structure was entirely rebuilt in our cryoEM map using coot (Waterhouse et al., 2018). Densities for the backbone and bases of ssDNA were clearly visible. Since the sequence of M13 ssDNA is different from one filament to another, a poly-dA (adenosine) DNA molecule was built in these averaged densities. Finally, ATPγS molecules were built in the corresponding densities of the map. *Sp*RecA model with ssDNA and ATPγS was refined against the cryoEM map using real-space refinement in Phenix (Phenix et al., 2010).

**Table 1:**
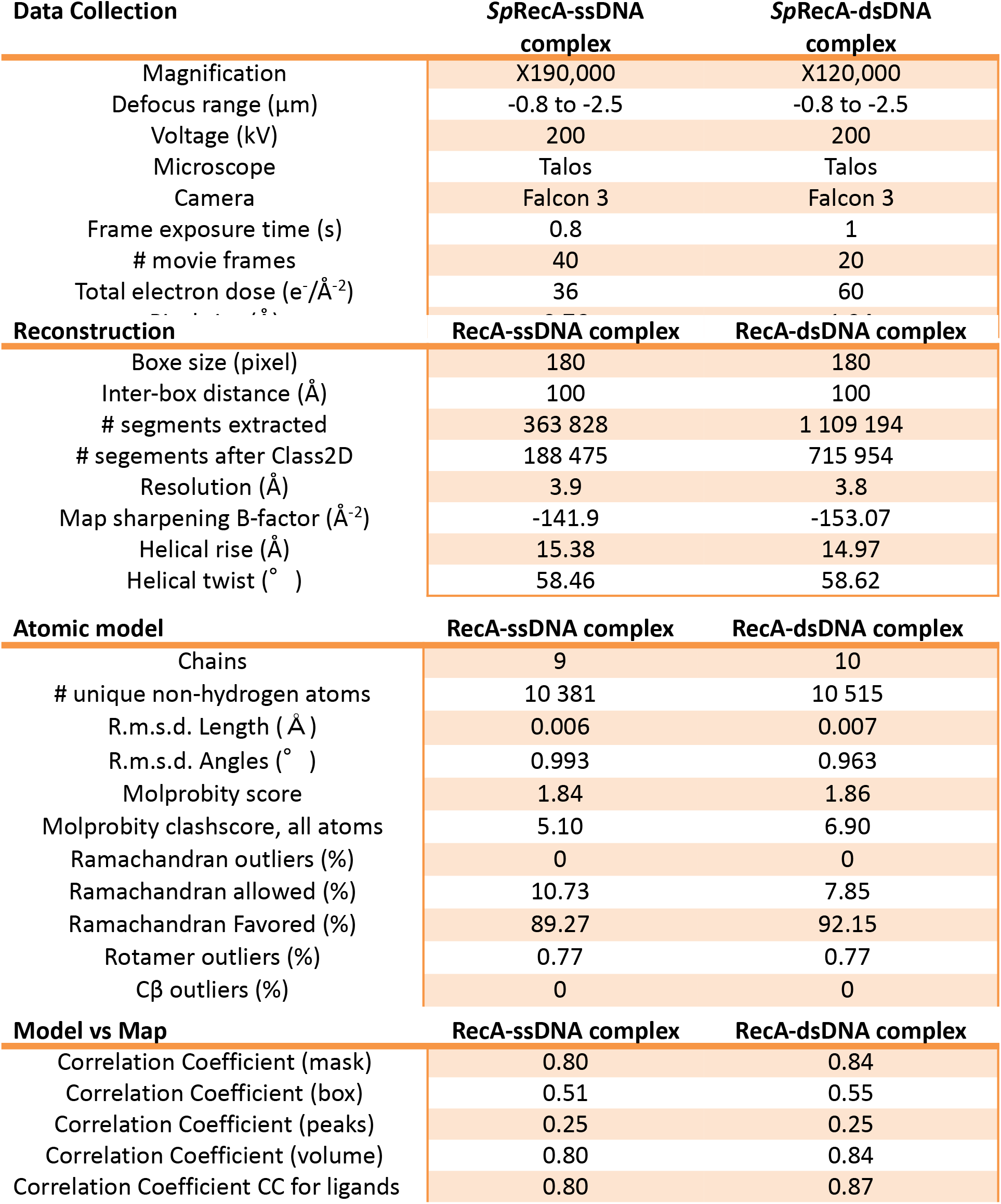
Cryo-EM structure determination and model tics for *Sp*RecA-ssDNA and *Sp*RecA-dsDNA complexes.

| Data Collection | *Sp*RecA-ssDNA complex | *Sp*RecA-dsDNA complex |
| --- | --- | --- |
| Magnification | X190,000 | X120,000 |
| Defocus range ($\mu\text{m}$) | -0.8 to -2.5 | -0.8 to -2.5 |
| Voltage (kV) | 200 | 200 |
| Microscope | Talos | Talos |
| Camera | Falcon 3 | Falcon 3 |
| Frame exposure time (s) | 0.8 | 1 |
| # movie frames | 40 | 20 |
| Total electron dose ($\text{e}^-/\text{\AA}^2$) | 36 | 60 |
| Reconstruction | RecA-ssDNA complex | RecA-dsDNA complex |
| Boxe size (pixel) | 180 | 180 |
| Inter-box distance ($\text{\AA}$) | 100 | 100 |
| # segments extracted | 363 828 | 1 109 194 |
| # segements after Class2D | 188 475 | 715 954 |
| Resolution ($\text{\AA}$) | 3.9 | 3.8 |
| Map sharpening B-factor ($\text{\AA}^{-2}$) | -141.9 | -153.07 |
| Helical rise ($\text{\AA}$) | 15.38 | 14.97 |
| Helical twist ($^\circ$) | 58.46 | 58.62 |
| Atomic model | RecA-ssDNA complex | RecA-dsDNA complex |
| Chains | 9 | 10 |
| # unique non-hydrogen atoms | 10 381 | 10 515 |
| R.m.s.d. Length ($\text{\AA}$) | 0.006 | 0.007 |
| R.m.s.d. Angles ($^\circ$) | 0.993 | 0.963 |
| Molprobity score | 1.84 | 1.86 |
| Molprobity clashscore, all atoms | 5.10 | 6.90 |
| Ramachandran outliers (%) | 0 | 0 |
| Ramachandran allowed (%) | 10.73 | 7.85 |
| Ramachandran Favored (%) | 89.27 | 92.15 |
| Rotamer outliers (%) | 0.77 | 0.77 |
| Cβ outliers (%) | 0 | 0 |
| Model vs Map | RecA-ssDNA complex | RecA-dsDNA complex |
| Correlation Coefficient (mask) | 0.80 | 0.84 |
| Correlation Coefficient (box) | 0.51 | 0.55 |
| Correlation Coefficient (peaks) | 0.25 | 0.25 |
| Correlation Coefficient (volume) | 0.80 | 0.84 |
| Correlation Coefficient CC for ligands | 0.80 | 0.87 |

The *Sp*RecA nucleofilament structure assembled and stabilized on ssDNA with ATPγS was found to be globally superimposable with the crystal structure of the *Ec*RecA-ssDNA nucleofilament (Figure 4a and 4c) obtained in presence of ADP-AlF_4_-Mg^2+^ (Chen et al., 2008). Each protomer binds to 3 nucleotides, organizing the ssDNA in triplets of a nearly B-form conformation that are separated from each other by 7.2 Å (Figure 4b and 4d).

**Figure 4:**
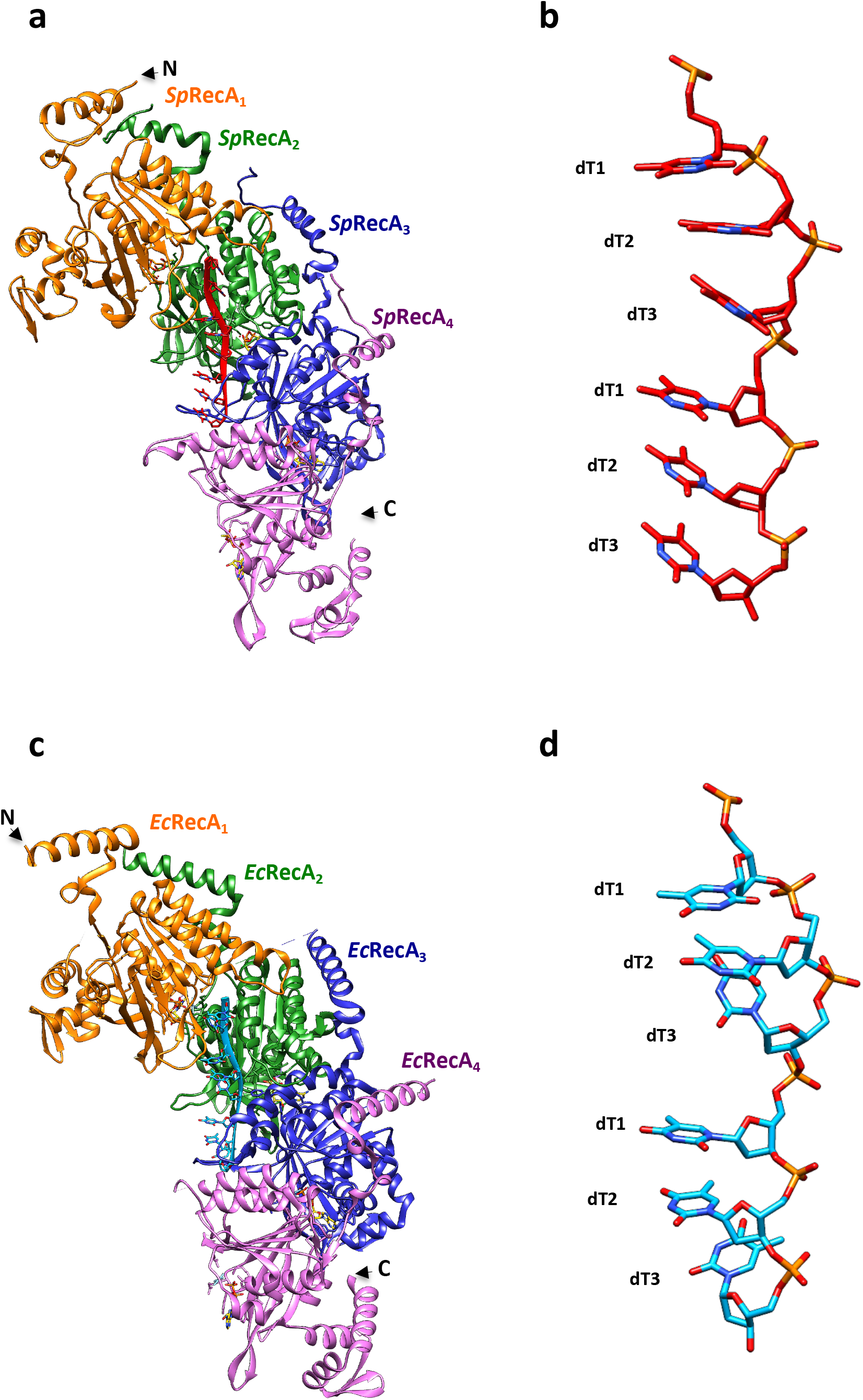
Structure comparison of the presynaptic nucleoprotein filaments from *Sp* and *Ec*. a and c. Structure of the RecA-ATPγS-dT complex from *Sp*RecA (a) and *Ec*RecA (c). Four RecA protomers are numbered from the N-terminus of the first protomer to the C-terminus of the last protomer, coloured in orange, green, blue and purple respectively. A single stranded DNA (ssDNA) molecule composed of 8 thymidine nucleotides bound to *Sp*RecA and *Ec*RecA are represented in red and blue respectively. Four ATPγS molecules are shown in gold. b and d. Zoom on the single strand B-form DNA from *S. pneumoniae* (b) and *E. coli* (d) presynaptic filaments. The ssDNA is numbered starting with the 5′-most nucleotide in each nucleotide triplet. The ssDNA binds with a stoichiometry of exactly three nucleotides per RecA, and the repeating unit of the DNA structure is a group of three nucleotides with a 3.5– 4.2□Å spacing.

The structure of each *Sp*RecA protomer in the nucleofilament appears to be very similar to the crystal structure of the *Ec*RecA protomer unbound to DNA 7/28/2022 4:39:00 AM. It is composed of a N-terminal extension (residues 9-55), a typical α/β ATPase core domain (residues 56-286) containing a canonical nucleotide binding motif and the conserved DNA interacting loops L1 and L2, and a globular C-terminal domain (residues 287-341). The charged C-terminal tail (residues 342 to 388) could not be resolved. In the *Sp*RecA filament, we numbered consecutive *Sp*RecA protomers along ssDNA in the 5’ end to the 3’ end direction. Within the nucleofilaments, the *Sp*RecA protomers interact mostly through their ATP binding domains. In addition, the N-terminal extension of the *Sp*RecA^n^ protomer lies on the ATPase domain of the adjacent *Sp*RecA^n-1^ protomer.

Within the *Sp*RecA nucleofilament, the DNA binding pockets are delineated by three consecutive *Sp*RecA protomers to accommodate a DNA triplet. In each pocket, the L1 and L2 loops of *Sp*RecA _n_ and *Sp*RecA^n+1^ protomers play a crucial role in contacting the ssDNA by encircling the DNA backbone (Figure5a). Residues from three consecutive *Sp*RecA protomers contribute to ssDNA binding through hydrogen bonds. Within the phosphate backbone of each triplet of nucleotides, from 5’ to 3’ end, the first phosphate group interacts with the backbone amide groups of E210 in *Sp*RecA^n^ and R226 in *Sp*RecA^n+1^, the second phosphate interacts with the backbone amide groups of G224 and G225 in *Sp*RecA^n+1^ and the third phosphate interacts with side chains from R209 in *Sp*RecA^n+1^ and S185 in *Sp*RecA^n+2^ (Figure 5b). The V212 residue found in the L2 loop inserts between consecutive triplets compensating the lack of base stacking in the inter-triplet junction (Figure 5b). The ATPγS binding pocket is shared by two consecutive *Sp*RecA protomers. In the *Sp*RecA^n^ protomer, the Walker A motif (residues 79-86) contacts ATPγS with conserved residues G84, K85 and T86 contacting the ATPγS phosphate groups. The third ATPγS phosphate makes several hydrogen bonds with the residues K265 and K267 in the RecA^n+1^, stabilizing the *Sp*RecA^n^ / *Sp*RecA^n+1^ interface. Finally, the highly conserved catalytic glutamate (E109) amongst RecA proteins is also found in the vicinity of the phosphate moieties.

**Figure 5:**
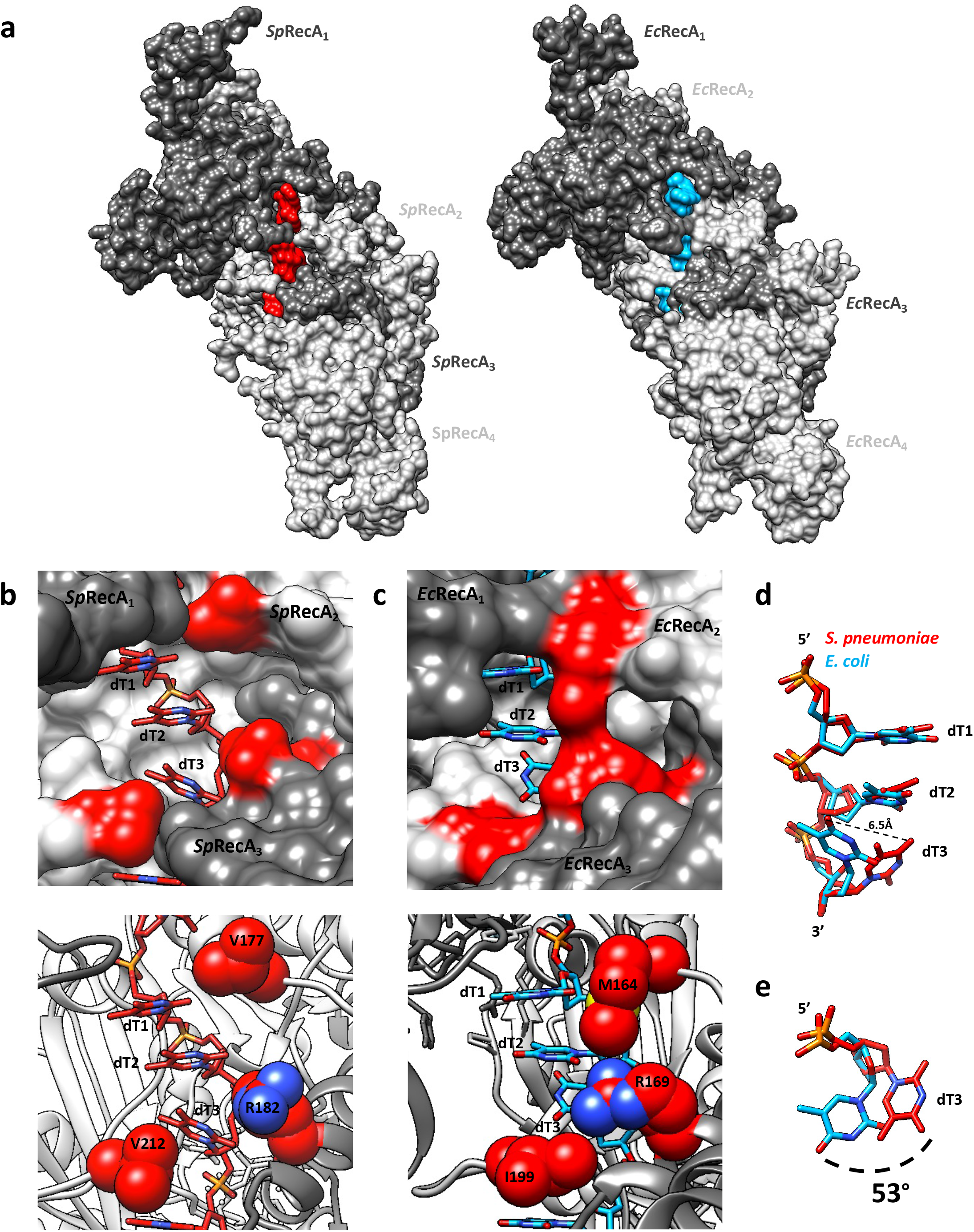
Interaction comparison between ssDNA and RecA protomers from from *Sp*and *Ec*. Surface representation comparaison of the RecA-ATPyS-dT complex from *S. pneumoniae* (*Sp*RecA) on the left and from *E. coli* (*Ec*RecA) on the right. Four RecA protomers are numbered from the N-terminus of the first protomer to the C-terminus of the last protomer, coloured in different grey. A single stranded DNA (ssDNA) molecule composed of 8 thimidine nucleotides bound to *Sp*RecA and *Ec*RecA are represented in red and blue respectively. b and c. Zoom of the RecA-ssDNA contacts from *S. pneumoniae* (b) and *E*. coli (c) presynaptic filaments. Each nucleotide triplet is bound by three consecutive RecA protomers. RecA protomers and ssDNA are numbered and coloured as the Figure 4.a. Residues V177, R182, and V212 from *S. pneumoniae* presynaptic filament (b) and residues M164, R169, and I199 from *E. coli* presynaptic filament (c) are coloured in red. d. Superimposition of the nucleotide triplets bound to *Sp*RecA (red) and *Ec*RecA (blue). The two first nucleotides can be superimposed while the last nucleotide of each triplet shows a difference in orientation, represented by a dotted line of 6.5Å long. e. Top view of the last nucleotide of each triplet superimposed and numbered dT3. The superimposition shows a shift of 53*°*.

The *Sp*RecA filament structure is in a native conformation, without protomeric fusion. Indeed, the published crystal structure of the pre- and post-synaptic *Ec*RecA filament was obtained using a chimera of six *Ec*RecA protomers truncated for Nter (1-30) and Cter (336– 353) residues and mutated to avoid oligomerization (C117M, S118V and Q119R). This chimera was bound to ssDNA and ADP-AlF_4_-Mg^2+^ (Chen et al., 2008). In *Sp*RecA filament, the presence of Nter or Cter regions does not modify the overall organization. When superimposed, *Ec*RecA and *Sp*RecA protomers have a RMSD of 1.392 Å. Similar helical parameters are found both for *Ec*RecA and *Sp*RecA presynaptic filaments. Structure-based alignment shows 55.86% identity between *Ec*RecA and *Sp*RecA sequences. Conservation of the residues is distributed across the whole structure. The ssDNA binding pocket and the L1/L2 loops are also particularly conserved between *Ec*RecA and *Sp*RecA. All interactions found between ssDNA and *Ec*RecA are also found in *Sp*RecA. The only notable differences are the V177 and V212 residues in *Sp*RecA, which correspond to the M164 and I199 residues in *Ec*RecA, respectively (Figure 5c). They are located at the tip of the L2 and L1 loops, respectively. These residues belonging to two consecutive RecA protomers close the L1/L2 loops around the primary ssDNA in the pre-synaptic nucleofilament and intercalate between the DNA triplet bound to RecA. However, one marked distinction stands out. Both recombinases stretch the ssDNA molecule the B-form of DNA in a non-uniform manner, and remarkably, while the third base of each triplet was found turned towards the interior of the *Ec*RecA protein filament, the three bases of the nucleotide triplet are all aligned toward the outer surface of the *Sp*RecA protein filament.

In parallel, we analyzed *Sp*RecA filamentation on dsDNA and, by contrast with the ssDNA matrix, we found that *Sp*RecA could self-assemble on dsDNA into long and stable filaments in the presence of ATPγS. We successfully analyzed their structure by cryoEM by applying a similar procedure as with filaments obtained on ssDNA. A 3.8 Å resolution map of these filaments has been obtained, in which we built and refined the structure of *Sp*RecA bound to dsDNA and ATPγS with a helical symmetry of 14.97 Å rise and 58.62 Å twist (corresponding to 6.14 subunits per turn). The overall structure of the individual *Sp*RecA protomer, as well as the interactions between *Sp*RecA protomers and with the ATPγS in this filament assembled on dsDNA are identical to those characterized for the filament assembled on ssDNA (Supplemental Figure 2b). Remarkably, however, the overall dsDNA B-form structure has been notably modified by polymerization of *Sp*RecA protomers. These were found to interact with one DNA strand as in the filament built with ssDNA. The complementary strand interacts with this primary DNA strand through Watson–Crick hydrogen bonds and makes very few contacts with *Sp*RecA protomers (Supplemental Figure 4). Interestingly, this structural organization of the dsDNA generated by *Sp*RecA polymerization appeared to be identical to the crystal structure of the dsDNA molecule resulting from the pairing of a ssDNA strand pre-bound by *Ec*RecA with its complementary ssDNA strand (Chen et al., 2008).

### Direct imaging of *Sp*RecA assembly on single molecules of DNA by TIRFm

Next, we undertook the analysis of nucleofilamentation dynamics of *Sp*RecA. To this end, we performed real-time observation of its polymerization on a single DNA molecule by Total Internal Reflection Fluorescence microscopy (TIRFm), following the same procedure previously developed for the study of *Ec*RecA nucleofilamentation (Bell et al., 2012). We used a DNA substrate composed of a central ssDNA gap of 8155 nucleotides flanked by biotinylated dsDNA ‘handles’ of 21080 and 24590 bases pairs (Figure 6a) and a fluorescently labeled *Sp*RecA (Alexa 488, named *Sp*RecA^A488^ hereafter) characterized for several activities. The purified labeled *Sp*RecA^A488^ was demonstrated to be active for ssDNA binding, D-loop formation and ssDNA-dependent ATP hydrolysis with a slight defect compared to the non-labeled protein (Supplemental Figure 5). DNA molecules were then attached to the surface of a streptavidin-coated glass coverslip in a microfluidic chamber and visualized by TIRFm. We detected interaction of *Sp*RecA^A488^ with DNA in the presence of ATPγS, but not with ATP, reproducing our previous TEM experiments (Figure 1d). *Sp*RecA filament formation first appeared as a single spot in the minute range.

**Figure 6.**
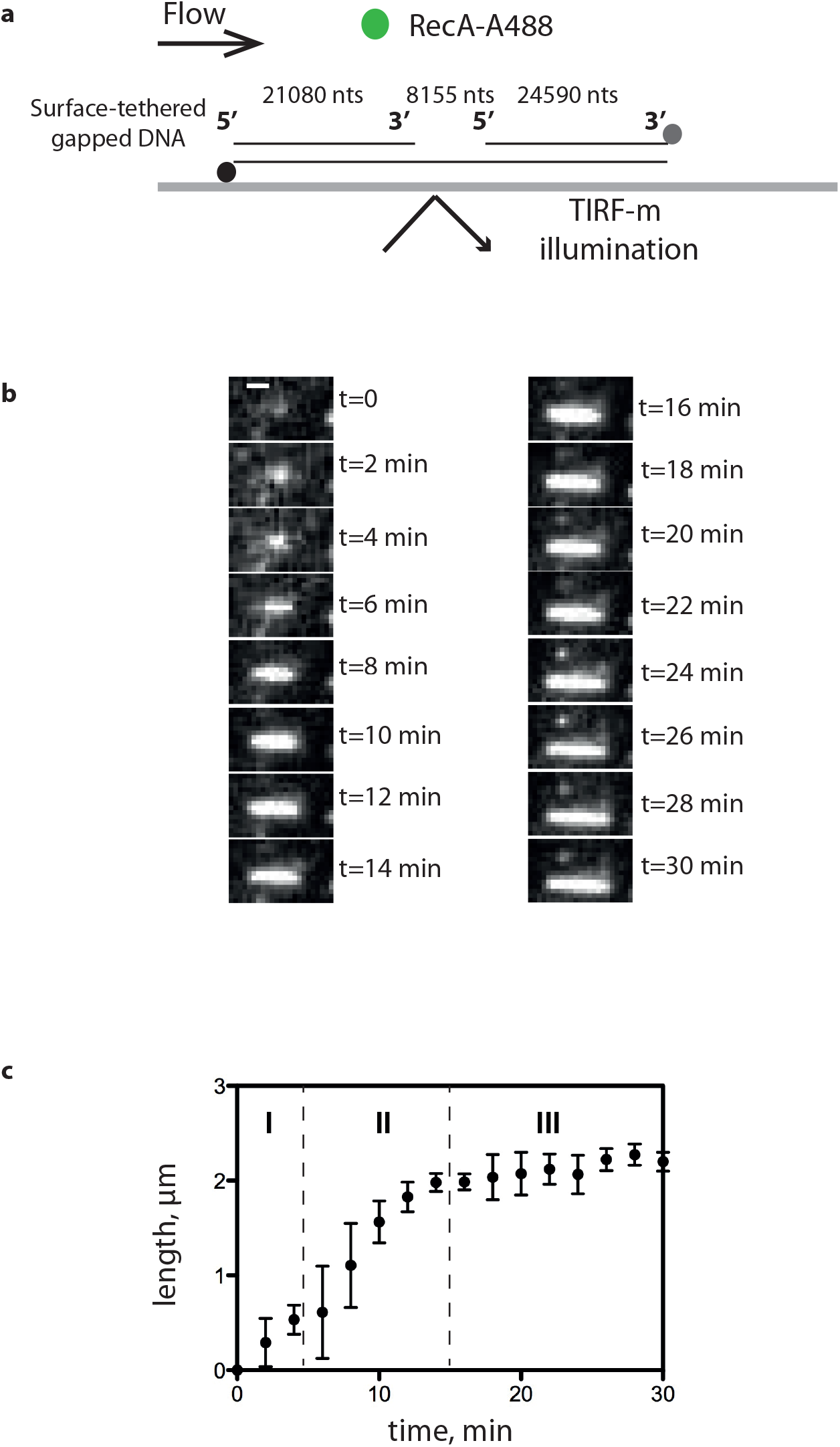
TIRFm analysis of *Sp*RecA assembly on single molecules of ssDNA. a. Schematic of the experimental set up combining TIRFm and microfluidics for direct imaging of *Sp*RecA^A488^ filament assembly in presence of ATPγS on a single molecule of ssDNA tethered within a microfluidic flow chamber. b. Sequential images of *Sp*RecA^A488^ filament assembly in presence of ATPγS. Scale bars represent 1µm and the time interval in minutes (min) is indicated in the images. c. The length of *Sp*RecA^A488^ filament clusters increases linearly with time. The plots are the average of 3 experiments and the standard error of the mean (sem) is represented.

Then, the size of individual *Sp*RecA^A488^ filaments gradually grew over time (Figure 6b) to reach a stable length after 15 min. The final filament occupies the place on the DNA molecule that is unbound by Sytox and, therefore, corresponds to the ssDNA portion. The averaged *Sp*RecA filament growth rate measured on 3 individual DNA molecules showed a consistent and reproducible polymerization into three distinct stages, referred to as initiation, elongation, termination (Figure 6c). The elongation rate of *Sp*RecA nucleofilament was 165 +/- 18 nm min^-1^, which is similar to the elongation rate previously reported measured for *Ec*RecA and measured on the same DNA molecule (50 to 500 nm min^-1^, Bell et al., 2012). The *Sp*RecA^A488^ filament assembly eventually reached a stable and maximum length of 2.1 +/- 0.1µm, which is 10 to 20-fold longer than *Sp*RecA nucleofilaments measured by TEM in the same experimental conditions (compare Figure 2c and Figure 3). A main difference between the TEM and TIRFm experiments is the application of a buffer flux in the microfluidic chamber in the later situation. Thus, it seems that the ssDNA stretching by the flow helps *Sp*RecA extension on longer distances, possibly by limiting the formation of ssDNA secondary structures that block its polymerization or stabilizing *Sp*RecA filament as it has been already observed for RAD51 (van Mameren et al., 2009). However, this measured length of the nucleofilament does not correspond to the maximum length of 4 µm expected for a saturating coverage of the ssDNA portion of the DNA substrate by one RecA molecule every 3 nucleotides (as demonstrated with the solved cryoEM structure of the *Sp*RecA nucleofilament; Figure 4).

Altogether, this TIRFm analysis shows that *Sp*RecA filament growth follows the same dynamics as that reported for *Ec*RecA. The two recombinases appear to differ in their intrinsic capacity to overcome some secondary structures to extend along the ssDNA matrix, but not in their elongation rate along ssDNA.

### *Sp*RecA ATP-dependent ssDNA binding mode

As we were not able to detect any *Sp*RecA assembly on DNA by TIRFm in the presence of ATP, we used Fluorescence Correlation Spectroscopy (FCS) and Fluorescence Anisotropy (FA) to measure the kinetics of formation of short *Sp*RecA polymers on ssDNA that could not have been detected by TIRF microscopy.

FCS allows the detection of fluorescently labeled molecules that diffuse through a sub-femtoliter detection volume, giving rise to intensity fluctuations in real time and at the millisecond scale (Figure 7a) and allowing to calculate their diffusion time within the observation volume *τ*_D_. We used ssDNA substrates with random sequences, labeled with Alexa 488 at the 5’ end. We tested several lengths of ssDNA substrates, i.e., 1000, 500 and 100 nucleotides long, and we were able to detect exploitable signal changes only for the small 100-mers. Upon addition of *Sp*RecA or *Ec*RecA and ATP, the diffusion time increased with time prior reaching a plateau that reports on the ssDNA assembly kinetics of the two recombinases (Figure 7b and 7c, for 250 nM and 400 nM of each RecA, respectively). Thus, in these conditions, we were able to detect *Sp*RecA and *Ec*RecA assembly on ssDNA in the presence of hydrolysable ATP and to compare their kinetics in those conditions. To this end, we measured the average half-time to reach the plateau value in each condition. This was slightly shorter for *Sp*RecA in both conditions, i.e. 72 sec for *Sp*RecA and 112 sec for *Ec*RecA at 250 nM, and 69 sec for *Sp*RecA and 132 sec for *Ec*RecA at 400 nM. In addition, the kinetics of assembly on ssDNA appeared to be clearly different for the two recombinases. Indeed, in the very early stage of *Ec*RecA assembly (Figure 7, blue curve, zoom), the curve showed a cooperative mode, whereas *Sp*RecA assembly was faster and showed no cooperativity.

**Figure 7.**
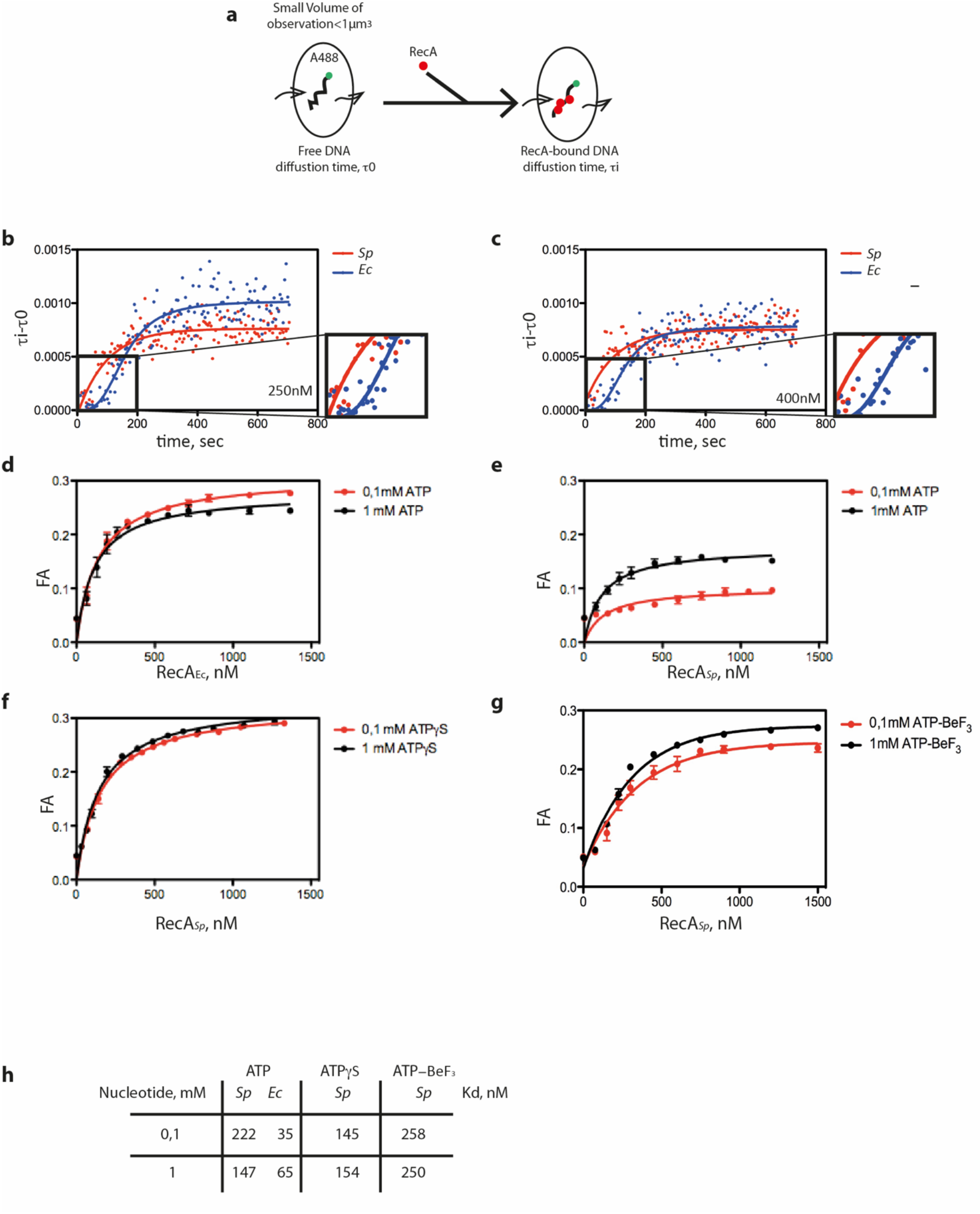
FCS analysis of *Sp*RecA and *Ec*RecA assembly on single molecules of ssDNA. a. Schematic of the experimental set up using FCS for the direct measurement of the change of diffusion time of fluorescently labeled (A488) ssDNA of 100 nucleotides length upon binding of *Sp*RecA and *Ec*RecA. b. and c. Averaged curve of 3 measurements of diffusion time at 250 nM of protein (b) showing a mean of half-time polymerization of 72,54 +/- 11,85 sec for *Sp*RecA and 111,99 +/- 18,24 sec for *Ec*RecA and at 400 nM of protein (c) showing a mean of half-time polymerization of 69+/- 6,0 sec for *Sp*RecA and 132, 35+/- 29,8 sec for *Ec*RecA. d-h. Equilibrium binding of *Ec*RecA (d.) of *Sp*RecA (e.) of *Ec*RecA in the presence of ATP. Fluorescence anisotropy (FA) variation with 1 mM ATP (black circles) and 0,1mM ATP (red circles) with a K_d_ of 65nM and 35nM in presence of 1 mM and 0,1 mM ATP, respectively. The plots are the average of 3 experiments and the standard error of the mean (sem) is represented. In the presence of ATPγS (f); ATP-BeF_3_ (g); Parameters table (K_d_) obtained from the above measurements (h).

In these FCS experiments, ATP hydrolysis by both recombinases triggered by their binding to ssDNA does not impact the stability of their interaction on ssDNA during such short periods of time. Complementarily, we characterized ssDNA binding affinity of *Sp*RecA and *Ec*RecA protein at steady state. To this end, we measured by fluorescence anisotropy (FA) their affinities constants for a short 65 nucleotides long fluorescent ssDNA molecule (T65). FA measurements were performed at 0.1 mM and 1 mM ATP, in large excess compared to the *Ec*RecA and *Sp*RecA concentrations used (Figure 7d and 7e, respectively). In those conditions, the measured apparent affinity for ssDNA (Kd) was 6 to 2-fold lower for *Sp*RecA than for *Ec*RecA at 0.1 mM ATP and 1 mM of ATP, respectively (Figure 7h). In addition, the maximum FA value reached ∼ 0.26 for *Ec*RecA, whereas it was less than 0.2 for *Sp*RecA, pointing at a different apparent molecular size of the nucleoprotein complexes. Thus, while the FCS analysis demonstrates that the two recombinases present a nearly equivalent half-time of association on ssDNA in the presence of ATP, the FA analysis indicates that they display a different ssDNA binding mode. Notably, no difference in the binding of *Ec*RecA to ssDNA was observed at the two ATP concentrations tested. In marked contrast, the plateau value reached for *Sp*RecA was found to be lower at 0.1 mM than at 1 mM ATP, and this latter value was lower than the one measured for *Ec*RecA. To test the impact of ATP hydrolysis in these differences, we reproduced these FA experiments in the presence of ATPγS or ATP-BeF_3_. Interestingly, in those conditions the ssDNA binding curves obtained for *Sp*RecA were found identical whatever nucleotide concentration used, either 0,1 or 1 mM and the value of the plateau matched with *Ec*RecA curves generated in the presence of ATP (Figures 7f and 7g to compare with 7d). This FA analysis showed that ATP hydrolysis modulates differently *Sp*RecA and *Ec*RecA interaction on ssDNA, despite both exhibit a similar ssDNA-induced ATP hydrolysis rate (Figure Supplemental 6). Altogether, these results show that *Sp*RecA binding on ssDNA appears markedly less stable upon ATP hydrolysis, pointing at a distinct and more dynamic mode of interaction with ssDNA for *Sp*RecA in comparison with that of *Ec*RecA. Also, ssDNA binding activities of both RecA proteins measured by FCS and FA revealed that ATP hydrolysis impacts differently the stability of their interaction on ssDNA, while they display a similar rate of ssDNA-dependent ATP hydrolysis.

### *Sp*RecA is more efficient than *Ec*RecA in a D-loop assay

Then, we measured the intrinsic ATP-dependent DNA strand-exchange activity of *Sp*RecA and *Ec*RecA. To this end, we used the D-loop assay illustrated in Figure 8b. In this assay, a 100-nucleotides (nts) linear oligonucleotide, fluorescently labeled with Cy3 at its 5’end, was incubated with increasing amounts of *Sp*RecA or *Ec*RecA and mixed with a homologous supercoiled plasmid. Following protein denaturation, the fluorescent D-loop product was separated from free ssDNA by agarose gel electrophoresis and quantified. *Sp*RecA was found to be up to three times more efficient than *Ec*RecA (3.1%+/-1.15 versus 1.29% +/- 0.57, respectively; Figure 1c). This result contrasts with a previous analysis reporting a less efficient HR activity of *Sp*RecA in comparison with *Ec*RecA (Grove et al., 2012). However, the HR assay used was markedly different. In this assay depicted in Figure 8a, the HR reaction is initiated by DNA strand exchange at one end of a linear dsDNA molecule with its complementary sequence on a long circular homologous ssDNA molecule (> 5000 nts) and is followed by DNA branch migration over a long distance to get the final product. By contrast, the D-loop product results from the invasive pairing between a short ssDNA molecule with its complementary sequence in a supercoiled dsDNA molecule. Thus, *Sp*RecA and *Ec*RecA appear to be oppositely and differently active in catalyzing the initial ssDNA pairing with a complementary sequence and in extending ssDNA recombination by DNA branch migration. Altogether, these functional divergences between these two bacterial RecA appear to stem from the different stability of their presynaptic filaments independent of the ATP hydrolysis rate.

**Figure 8.**
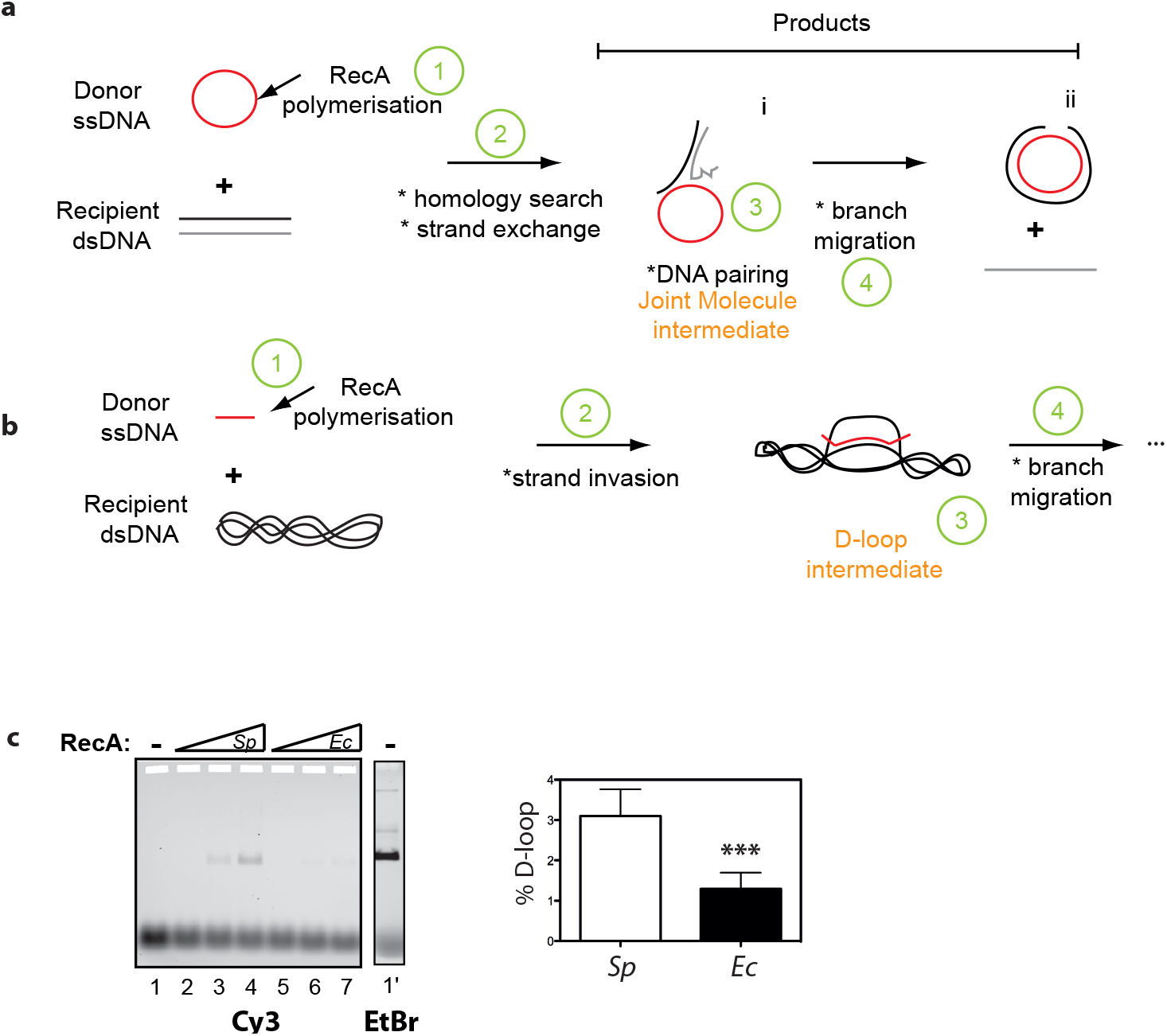
Comparison of *Sp*RecA and *Ec*RecA recombination activity in a D-loop assay. a and b. Schematics of *in vitro* DNA strands exchange assays commonly used to measure the recombination activity of HR recombinases; the D-loop assay is depicted in b. c. Left: deproteinized agarose gel of the D-loop reaction performed in presence of 10 nM of cy3 oligonucleotide (100 mers) 5 nM (1 ml) of pUC18 vector, and increased amount (150 to 600 nM) of *Sp*RecA or *Ec*RecA as indicated above the gel (*Sp* and *Ec*, respectively). Right: quantification of the D-loop product generated at 600 nM of *Sp*RecA and *Ec*RecA concentration. The percentage of D-loop formed is given as mean values +/- standard error of the mean (sem) of three reactions.

## Discussion

We report a comprehensive molecular study of *Sp*RecA functional properties, which provides important insights into its DNA interaction properties in relation with its DNA strand exchange activities. This *in vitro* analysis shows that *Sp*RecA markedly differs from the *Ec*RecA paradigm in the early stages of HR. Our findings collectively concur to the conclusion that the main deviation between the two HR recombinases mostly stems from their ATP-dependent ssDNA binding mode and independently of their ATPase rates. Our studies also highlight the lack of synergy between *Sp*RecA and its two cognate and paralogous SSB proteins in elongating its presynaptic filamentation, a defect revealed to be compensated by the conserved RadA helicase, previously known to act coordinately with RecA on postsynaptic HR intermediates. In addition, they support a model of HR mechanism in which the *Sp*RecA presynaptic filament would be more efficient than *Ec*RecA in homology search and ssDNA pairing within a recipient complementary dsDNA molecule.

### Key variations in the ATP-dependent ssDNA interaction dynamics of *Sp*RecA and*Ec*RecA

A central intermediate of the HR mechanism is the presynaptic filament, which is dynamically assembled and disassembled on ssDNA by ATP binding and hydrolysis between the protomers of the recombinase (Liu et al., 2011). This ssDNA-dependent ATP cycle is not uniformly conserved between bacterial RecA, leading to various lengths of presynaptic filaments (Cox, 2007; Morrical, 2015). We report here that *Sp*RecA interaction with ssDNA in the presence of ATP evaluated by FA analysis is more dynamic than that of *Ec*RecA (Figure 7). This finding indicates that *Sp*RecA forms shorter nucleofilaments than *Ec*RecA, as further supported by TEM analysis (Figure 2c and 2g, respectively). However, this marked difference between the two recombinases is not due to a different affinity for ATP, nor to a different ssDNA-dependent ATP hydrolysis rate, nor to a different kinetic in ATP-dependent interaction with ssDNA (Supplemental figure 1). Thus, a possible cause of the limited extension of the presynaptic filament of *Sp*RecA would be a lower binding stability of its protomers on ssDNA. Within the HR presynaptic filament, each protomer interacting with ssDNA is further stabilized *via* interaction with two adjacent protomers through ATP binding. Upon ATP hydrolysis, protomers located at the tips of the filament are less stably bound to ssDNA, as they are engaged in only one interaction with an adjacent protomer. Thus, it has been shown by biochemical and SM analysis that the *Ec*RecA filament mainly disassembles at the 5’ side and grows in the 3’ direction of the ssDNA (Bell and Kowalczykowski, 2016b). In direct line with such a polymerization dynamic, the 5’ terminal protomer of the *Sp*RecA filament might be less stable on ssDNA upon ATP hydrolysis than in the case of the *Ec*RecA filament. In addition, the release of Pi from the ADP-Pi product will change the interactions between the two protomers, which will alter their interaction with ssDNA. Thus, a possible source of difference between *Sp*RecA and *Ec*RecA impacting the length of their presynaptic filaments would be the ADP.Pi release and/or ATP turnover at the interface of two protomers bound to ssDNA.

Furthermore, even under stabilizing conditions that restrain ATP hydrolysis (with the use of ATPγS or by adding BeF3 to ATP), *Sp*RecA appears less prone than *Ec*RecA to elongate on ssDNA (Figure2). We interpret this difference as a lower capability of *Sp*RecA to melt ssDNA secondary structures in comparison with *Ec*RecA, limiting differentially their filament growth. The longer *Sp*RecA presynaptic filaments observed in TIRFm than in TEM experiments supports this proposal (Figs2 and3). SM observation of *Sp*RecA filamentation by TIRFm is performed in real time in a microfluidic chamber on an attached DNA molecule and under flux, which will physically extend the ssDNA and limit its self-pairing into secondary structures. As a result, *Sp*RecA could extend its polymerization, stabilized by limiting ATP hydrolysis, on a longer distance than on an unstretched ssDNA molecule as in the TEM experiments.

Another distinct ssDNA interaction property between these two recombinases has been uncovered from the characterization by cryoEM of the *Sp*RecA nucleofilament structure stabilized by ATPγS. This structure is superimposed to a large extent on the *Ec*RecA filament resolved by crystallization (Chen et al., 2008). In both *Sp*RecA and *Ec*RecA nucleofilaments, the ssDNA molecule is bound into an identical helical and extended conformation organized in triplets of nucleotides. However, the 3 bases of each triplet of nucleotides are fully exposed toward the exterior in the *Sp*RecA filament, contrasting with the *Ec*RecA filament where the third base of each nucleotide triplet is flipped inward (Chen et al., 2008). These characteristics suggest that the ssDNA conformation in the *Sp*RecA filament is potentially more favorable for the homology search.

### Lack of SSB assistance in the extension of *Sp*RecA presynaptic filamentation

One of the key roles of SSB in the early HR steps is to assist presynaptic filament extension by melting out ssDNA secondary structures (Bianco, 2017). This interplay originally characterized between RecA and SSB of *E. coli* has been generalized to RecA of many other species, with the marked exception of *S. pneumoniae*. Indeed, either of the two pneumococcal SsbA and SsbB proteins, or *Ec*SSB were found to inhibit *Sp*RecA binding to ssDNA, as deduced from their inhibition of the ssDNA dependent *Sp*RecA ATPase activity (Grove and Bryant, 2006). Here, we directly observed by TEM analysis that any of these SSB outcompetes ATP-dependent *Sp*RecA polymerization on long ssDNA molecules. Furthermore, their addition to the short *Sp*RecA filaments stabilized by ATP γS simply conduct to their binding on ssDNA portions unbound by *Sp*RecA, without promoting the extension of *Sp*RecA nucleofilaments as in the case of *Ec*RecA (Figure 2). This result firmly demonstrated the lack of assistance by any SSB in elongating *Sp*RecA polymers on ssDNA. They also indicate that the inhibition by SSB proteins of the ATP-dependent *Sp*RecA interaction on ssDNA is the result of a more stable binding of SSB in comparison to the highly dynamic binding of *Sp*RecA, leading to a full occupancy of the ssDNA by SSB in these conditions. This also shows that SSB can bind to ssDNA parts that are inaccessible to *Sp*RecA and inferred to be secondary structures.

An elegant genetic screen of *Ec*RecA mutants more efficient in conjugational recombination resulted in the selection of several point mutants that were all found to exhibit *in vitro* a greater persistence on ssDNA and a more efficient displacement of SSB than wild type *Ec*RecA (Kim et al., 2015). The *Ec*RecA region randomly mutated in this screening corresponds to the large N-ter region involved in RecA subunit-subunit interaction. This shows that modulations in this interacting interface could impact the intrinsic ssDNA interacting and polymerizing property of RecA on ssDNA. However, comparison of this interaction surface between *Sp*RecA and *Ec*RecA could not highlight a particular difference that would explain the lower persistence of *Sp*RecA on ssDNA that we report here. In addition, other subtle variations between RecA proteins might influence their intrinsic stability on ssDNA. Indeed, another possible source of variation could be the residues engaged in direct interaction with ssDNA, as we found here in the structure of the presynaptic filaments of *Sp*RecA in comparison with *Ec*RecA (see above). However, further studies are needed to establish whether this different organization of the presynaptic filament modifies their dynamism.

### An unprecedented role of the RadA HR effector in extending RecA presynaptic filamentation

The less stable ssDNA binding in *Sp*RecA filament leads to their limited extension, which is impeded by SSB proteins or ssDNA secondary structures and unfavorable for the branch migration step in HR reaction (Grove and Bryant, 2006). SSB proteins are well known effectors that assist RecA dynamics and filament length (Roy et al., 2009). For *E. coli, Pseudomonas aeruginosa, Neisseria gonorraheae, Herbaspirillum seropedicae* or *Bacillus subtilis* (*Bs*) RecA proteins, SSB proteins remove structures in ssDNA to facilitate formation of *Ec*RecA nucleoprotein filaments on ssDNA (Gruenig et al., 2010). In the experimental conditions tested here, *Sp*SsbA or *Sp*SsbB protein improve only very slightly or compete with the *Sp*RecA filament extension. Like for *Dr*RecA, *Sp*RecA ATP hydrolysis is inhibited by *Sp*SsbA or *Ec*SSB. So, regarding SSB proteins, *Sp*RecA showed a distinct behavior shared with *Dr*RecA. In contrast, *Sp*RadA helicase enhanced the *Sp*RecA filament extension. It does so without co-polymerizing with it. The use of the ATP hydrolysis mutant of *Sp*RadA (*Sp*RadA^K101A^) showed that the ATP hydrolysis activity of *Sp*RadA is required to enhance *Sp*RecA filament extension. This strongly suggests that helicase activity of *Sp*RadA could remove ssDNA secondary structures to help *Sp*RecA extension. Interestingly, RecA filament growth is well known to proceeds from 5’ to 3’ on ssDNA, which is also the translocation directionality of *Sp*RadA when acting as helicase (Marie et al., 2017).Thus, *Sp*RadA not only acts in HR mechanism at the post-synaptic step by promoting DNA branch migration (Cooper and Lovett, 2016; Marie et al., 2017; Torres et al., 2019) but also at the presynaptic step, by relieving the stem-loop structures that form on ssDNA and that impede RecA polymerization. Interestingly, *Sp*RecA is markedly inefficient in directing these two HR steps by itself (this study; (Grove and Bryant, 2006)). By marked contrast, we found that SpRecA is intrinsically highly efficient in promoting homologous ssDNA pairing in dsDNA template, even more than the *Ec*RecA paradigm (Figure 8c).

Altogether, this detailed structural and biochemical analysis of ATP-dependent DNA interacting properties of *Sp*RecA points at their balanced intrinsic efficiency by comparison with the *Ec*RecA paradigm. Also, these activities could be differently compensated by accessory effectors. These key variations on the conserved RecA-directed HR mechanism points at its adaptation amongst bacterial species, which could reflect specific needs and/or its particular integration with other processes at work on their genome.

## Material and methods

### Proteins

*Sp*RecA was purified as previously described (Marie et al., 2017). The *Ec*RecA protein was purified in a similar manner and then compared to commercial *Ec*RecA (NEB). The protein activity of commercial NEB and purified proteins was equivalent, so we used for this study either commercial or purified *Ec*RecA.

### D-loop assay

The basic reaction solution contained 10 mM Tris-Cl (pH 7,5), 0,1 mg/ml BSA, 8 % glycerol, 0,5 mM DTT or TCEP, 50 mM NaCl, 10 mM MgAc, 2 mM ATP, 10 nM of 5’ Cy3 100-mer oligonucleotide (5’-TGCTTCCGGCTCGTATGTTGTGTGGAATTGTGAGCGGATAACAATTTCACACAGGAAACAGCTATGACCATGATTACGAATTCGAGCTCGGTACCCGGGG-3’) homologous to pUC18 sequence, and RecA (150 to 600 nM). After incubation of RecA with the ssDNA (oligonucleotide) for 10 min at 37 °C, we added 5 nM of pUC18 vector into the reaction and further incubated 10 min at 37°C to allow oligonucleotide-pUC18 pairing (D-loop). The reaction was then kept on ice. The reaction was quenched (or deproteinized) with 1% SDS / 10 mM EDTA (final concentrations). 0,5 l of loading buffer (Xylene cyanol in 30% glycerol) was added and reactions analysed by electrophoresis on a 1,2 % agarose gel in a Tris-Acetate-EDTA buffer at room temperature, 6 V/cm for 1h in order to identify and estimate properly the amount of D-loop created. We detected the free and the bound Cy3 labelled oligonucleotides by a Fluor imager (Typhoon trio-Fuji-GE-healthcare) with an Abs/Em of 532/580 nm. Quantification of the proportion of D-loop created in this assay was performed with the Multigauge and Excel softwares.

### TEM analysis

For transmission electron microscopy studies, a fraction of the filament formation reactions described above was diluted and handled as previously described (Dupaigne et al., 2008). For statistical analysis of the length of filaments, 30 to 50 molecules were analyzed for each reaction. For RecA filament formation, 15 µM (nucleotides) ΦX ssDNA were first incubated with 5 µM *Sp*RecA or *Ec*RecA 3 minutes at 37°C in a buffer containing 10 mM Tris-HCl pH 7,5, 50 mM KCl, 5 mM MgCl_2_, 1 mM DTT and either 1,5 mM ATP or 1 mM ATPgammaS or 1,5 mM ATP plus 1,5 mM BeF_3_.

### Fluorescence labeling

*Sp*RecA^A488^ was made by covalently modifying primary amines (lysines or N-ter) of the protein with Alexa 488-succinidimyl ester (Molecular Probes, ThermoFisher), in presence of an excess of ssDNA (M13 mp18, NEB) and ATPγS (Roche) in order to preserve both ATP and ssDNA binding surfaces of the purified recombinant protein. The free fluorescent probe was removed by a step of gel filtration chromatography (Superdex 200 Increase 10/300 GL; GE Healthcare) in the reaction buffer 50mM Tris (pH7.5), 300mM NaCl, 1mM DTT. To remove any ATP or DNA contaminant, a final step of Anion Exchange chromatography was performed (MonoQ column GE Healthcare). RecA^f^ was prepared as previously described [20]. The ssDNA binding activity of RecA was determined by monitoring the ATP hydrolysis rate of RecA at increasing concentrations of ATP.

### Production of DNA substrates

Gapped DNA substrates were prepared as described previously (Bell et al., 2012). The short fluorescent ssDNA substrate used in FCS experiments was prepared with synthetic oligonucleotides (Eurogentec) labeled either with Biotin or Alexa-488 in 5’ in order to generate a Biotin-labeled DNA strand and a fluorescently-labeled DNA strand (Sequence : Biotin-5’GCTTGCATGCCTGCAGGTCG3’; Alexa488-5’GCGGATAACAATTTCACACAGG3’) by PCR-amplification using the pUC18 plasmid as template. After PCR amplification (Volume =2 ml), the PCR reactions were loaded on Hi-Trap Streptavidin column (GE Healthcare). By addition of 60 mM NaOH, the fluorescent DNA strand is eluted, while the Biotin DNA strand retains on the column. The fluorescent ssDNA is then precipitated by Chloroform/Isopropyl alcohol, resuspended in 10 mM Tris-HCl pH 7.5; 50 mM NaCl and qunatified using a Nanodrop spectrophotometer.

### Direct Imaging of RecA assembly on single molecules of ssDNA

A gapped ssDNA substrate was prepared and biotinylated as described in (Bell et al., 2012) The gapped ssDNA molecules were injected into a flow cell and tethered to the surface of a coverslip via biotin-streptavidin interactions. Flow cells (4 mm × 0.4 mm × 0.07 mm) were assembled using a glass slide, a coverslip, and double-sided tape (3M Adhesive Transfer Tape 9437). Ports were drilled into the glass microscope slide, and flow was controlled using a motor-driven syringe pump (Amitani et al., 2010; Forget and Kowalczykowski, 2012). The surface of the coverslip was cleaned by the subsequent injection of 1 M NaOH for 10 min, rinsed with water and equilibrated in buffer containing 20 mM TrisOAc (pH 8.0), 20% sucrose and 50 mM DTT. The surface was then functionalized by injecting the above buffer containing 2mg/ml biotin-BSA (Pierce) and incubated for 10 min, rinsed with buffer, equilibrated with 0.2 mg/ml streptavidin (Promega) for 10 min and then blocked with 1.5 mg/ml Roche Blocking Reagent (Roche) for 10 min. For imaging, the gapped DNA were allowed to incubate in the flow cell in the absence of flow for approximately 5–15 min.

### FCS measurements

Fluorescence correlation spectroscopy was performed on a custom-built setup with Pulse Interleaved Excitation (PIE) and Time Correlated Single Photon Counting (TCSPC) detection as described elsewhere (Olofsson and Margeat, 2013). The FCS measurements were performed in the presence of the indicated amount of RecA proteins, 5 nM fluorescently labeled ssDNA (fluorescent probe: Alexa-488; size: 100 bases), in a buffer containing10 mM Tris-HCl pH7.5; BSA 0,5 mg/ml; 4 mM MgCl_2_; 50 mM NaCl; 0,5 mM DTT.

### Equilibrium Anisotropy fluorescent binding assays

Titrations to monitor the binding of RecA to ssDNA were performed by monitoring the anisotropy of fluorescence enhancement at 25°C, using a Horiba fluorescence spectrophotometer set at an excitation wavelength of 495 nm and an emission wavelength of 520 nm. Excitation and emission slits were set to a bandwidth of 10 nm. Titrations were performed in 25 mM Tris-HCl (pH 7.5), 1 mM DTT, 25mM NaCl, 2.5% glycerol, 10mM Mg Cl_2_ and the indicated concentration of nucleotide. The Anisotropy of fluorescence values were corrected for dilution. An increased amount of RecA was added to the reaction solution containing the 25nM of polydT of 65-mers. Data fitting using One-site-specific binding model was performed using GraphPad Prism. All equilibrium titrations were performed 3 times and the curves shown are the average of three with SEM represented.

### Presynaptic and postsynaptic complex assembling

All the reaction steps were carried out at 37 °C. For assembling presynaptic filaments, □X174 Virion single strand DNA (New England BioLabs) at 10 ATPγS g.mL^-1^ was incubated with RadA at 50 µg.mL^-1^ for 1 min in the reaction buffer comprising of 10 mM HEPES pH 7.5, 100 mM NaCl, 50 mM KCl, 0.5 mM DTT and 1.5 mM ATPγS; 50 µM Mg Cl_2_. Then, *Sp*RecAwas added at final concentration of 200 µg.mL^-1^ for 3.5 h at 37°C. For assembling postsynaptic filaments, Lambda double strand DNA (New England BioLabs) at 10 □g.mL^-1^ was incubated with *Sp*RecA at 200 µg.mL^-1^ for 3.5 h at 37 °C in the same reaction buffer. Complex formation was checked by negative stain on a CM120 electron microscope (FEI/Thermo Ficher).

### Cryo-EM specimen preparation and electron microscopy data acquisition

For cryo-EM analyses, 3.5 µl of sample were deposited on glow-discharged Lacey carbon grids, blotted with filter paper to remove excess sample for 4 s, and plunge-frozen in liquid ethane using a FEI Vitrobot Mark IV (FEI/Thermo Ficher) with a blotting force of 0 in an environment with 100% humidity and 4 °C temperature. Cryo-EM images were acquired on a Falcon 3 direct detector in counting mode for the presynaptic complex and in linear mode for the postsynaptic complex on a FEI Talos Arctica at 200 kV. For the presynaptic complex, a magnification of 190,000 x was applied to record 40 movie frames with an exposure time of 0.8 s using a dose rate of 0.9 electrons per Å^2^ per frame for a total accumulated dose of 36 electrons per Å^2^ at a pixel size of 0.76 Å. For the postsynaptic complex, a magnification of 120,000 x was applied to record 20 movie frames with an exposure time of 1 s using a dose rate of 3 electrons per Å^2^ per frame, resulting in a total accumulated dose of 60 electrons per Å^2^ at a pixel size of 1.24 Å. The final datasets were composed of 2896 (for the presynaptic complex) and 2364 (for the postsynaptic complex) micrographs with defocus values ranging from -0.8 to -2.5 µm.

### Helical reconstruction

Similar procedures were applied to the presynaptic complex and the postsynaptic complex datasets using helical reconstruction methods in RELION 2.1 (S and Shw, 2017). All frames were corrected for gain reference, binned by a factor of 2 only for the presynaptic complex, motion-corrected and dose-weighted using MOTIONCOR2 (Zheng et al., 2017) Contrast transfer function (CTF) parameters were estimated by CTFind-4.1 (Rohou and Grigorieff, 2015).

Particles on micrographs of the presynaptic complex were picked manually in box sizes of 180 pixels and with an inter-box distance of 100 Å. Then, picked particles were classified into two-dimensional class averages to identify homogeneous subsets using a regularization value of *T*=2. Selected classes were used as references for autopicking in RELION 2.1 (Scheres, 2012). The total number of initial extracted segments (25,653) was reduced to 7,254 by subsequent rounds of two-dimensional classifications. After the best two-dimensional classes were selected, a first three-dimensional reconstruction was done using featureless cylinder of 125 Å in diameter as an initial model (Chen et al., 2008). This was achieved by refining without imposing any helical symmetry and. This yielded a map at 7.6Å in which helical symmetry was already apparent.

Then, this map was used as reference for new autopicking on the micrographs of both complexes. The total number of extracted particles (363,828 segments for the presynaptic and 1,109,194 segments for the postsynaptic) was reduced to 188,475 and 715,954 by subsequent rounds of two-dimensional classifications. High-resolution refinements were performed in RELION’s 3D auto-refinement using the non-symmetrized map as a reference, optimizing both the helical twist (58.46° and 58.62° respectively) and rise (15.38 Å and 14.97 Å respectively) (S and Shw, 2017). The final resolution was 3.9 A□ for the presynaptic complex and 3.8 Å for the postsynaptic complex, calculated with two masked half-maps refined independently, according to the gold standard Fourier shell correlation (FSC) 0.143 criterion using RELION. Local resolution, calculated with RELION with a B-factor applied of −141.9 and -153.07 respectively, retrieved a range between 3.7 and 7.7 A□. All of the densities obtained were subjected to Auto-sharpening (Afonine et al., 2018) in the Phenix software package.

### Model building and refinement

The initial atomic model of *Sp*RecA protomers in both presynaptic and post-synaptic complexes were generated from the crystal structure of *E*.*coli* RecA (PDB ID: 3cmw and 3cmx) by SWISS-MODEL (Schwede et al., 2003). Rigid-bodies, comprising four molecules of ATPγS and the DNA, were docked into the autosharpened electron density map in UCSF-Chimera (Pettersen et al., 2004). The coordinates of the obtained single-chain model were modified manually using Coot and refined with repeated rounds of Phenix real-space refine function. The structure was further refined in real-space in PHENIX with secondary structure restraint (Adams et al., 2010). The atomic models were validated using the Cryo-EM validation tools of Phenix (Afonine et al., 2018). Briefly, each model was firstly refined against the sharpened map (Supplemental data). To monitor the refinement of the model and avoid over-fitting, the final model was refined against one half map and tested against the other half map by calculating the Fourier Shell Correlation curves (not reported), which indicated that the refinement of the atomic coordinates did not suffer from over-fitting.

## ACKOWLEDGMENTS

This work was funded by: the Centre National de la Recherche Scientifique, University Paul Sabatier; the Agence Nationale de la Recherche (grant ANR-10-BLAN-1331); the IdeX Toulouse funding for Emergence with the SMART project ‘Single Molecule Analysis of Homologous Recombination’ attributed to MH and for Equipment with the ‘Go ahead in life sciences in Toulouse’ project; the Fondation de la Recherche Médicale (FRM N° ING20150532556) for the salary of SS; the European Research Council (ERC) consolidator grant TransfoPneumo attributed to RF; the France Bio Imaging (FBI) funding attributed to EM & MH. We thank Chantal Prevost for helpful discussions

