## Supplemental Figures for "Assembly mechanism and cryoEM structure of RecA recombination nucleofilaments from *Streptococcus pneumoniae*"

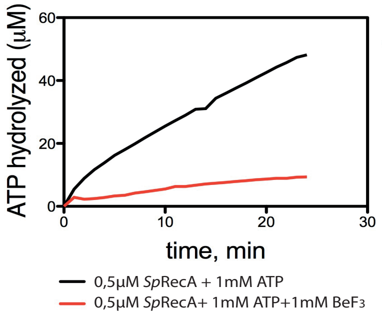


**Supplemental Figure1:** Comparisons of ATP hydrolyzed rates for 0,5 µM of proteins *Sp*RecA/ *Sp*RecA $^{A488}$ in presence or not of $BeF_3$.

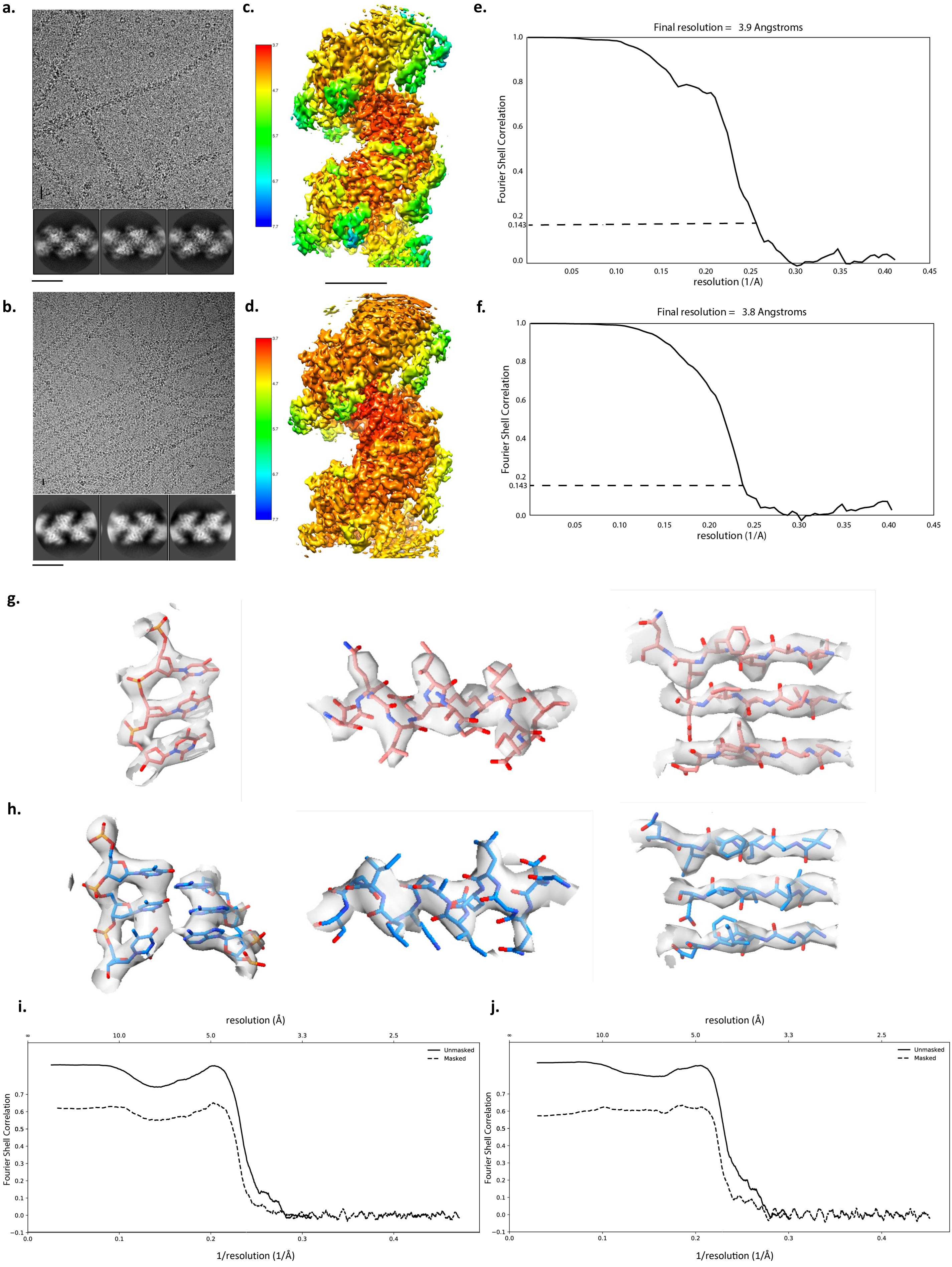

a.
b.
c.
d.
e.
Final resolution = 3.9 Angstroms
Fourier Shell Correlation
resolution (1/A)
f.
Final resolution = 3.8 Angstroms
Fourier Shell Correlation
resolution (1/A)
g.
h.
i.
resolution (Å)
Unmasked
Masked
Fourier Shell Correlation
1/resolution (1/Å)
j.
resolution (Å)
Unmasked
Masked
Fourier Shell Correlation
1/resolution (1/Å)

**Supplemental Figure 2. Analysis of model quality and refinement.**

a and b. Representative cryo-micrograph of RecA-ATPγS-ssDNA (a) and RecA-ATPγS-dsDNA (b) complexes from *S.pneumoniae* at the top (scale bar=200 Å), with typical 2D classes of the filament at the bottom (scale bar=100 Å). c and d. Surface representation with the local resolution of RecA-ATPyS-ssDNA (c) and RecA-ATPyS-dsDNA (d) (scale bar=50 Å). e and f. Fourier shell correlation (FSC) of the final reconstruction of RecA-ATPyS-ssDNA (e) and RecA-ATPyS-dsDNA (f). The resolution limit was calculated at the cut-off 0.143. g. Representative local density of a triplet (left), an !-helix (middle) and a β-strand (right) of RecA-ATPyS-ssDNA. f. Representative local density of a double triplet (left), an !-helix (middle) and a β-strand (right) of RecA-ATPyS-dsDNA. i and j. Fourier Shell Correlation curve calculated for the map versus model of RecA-ATPyS-ssDNA (i) and RecA-ATPyS-dsDNA (j).

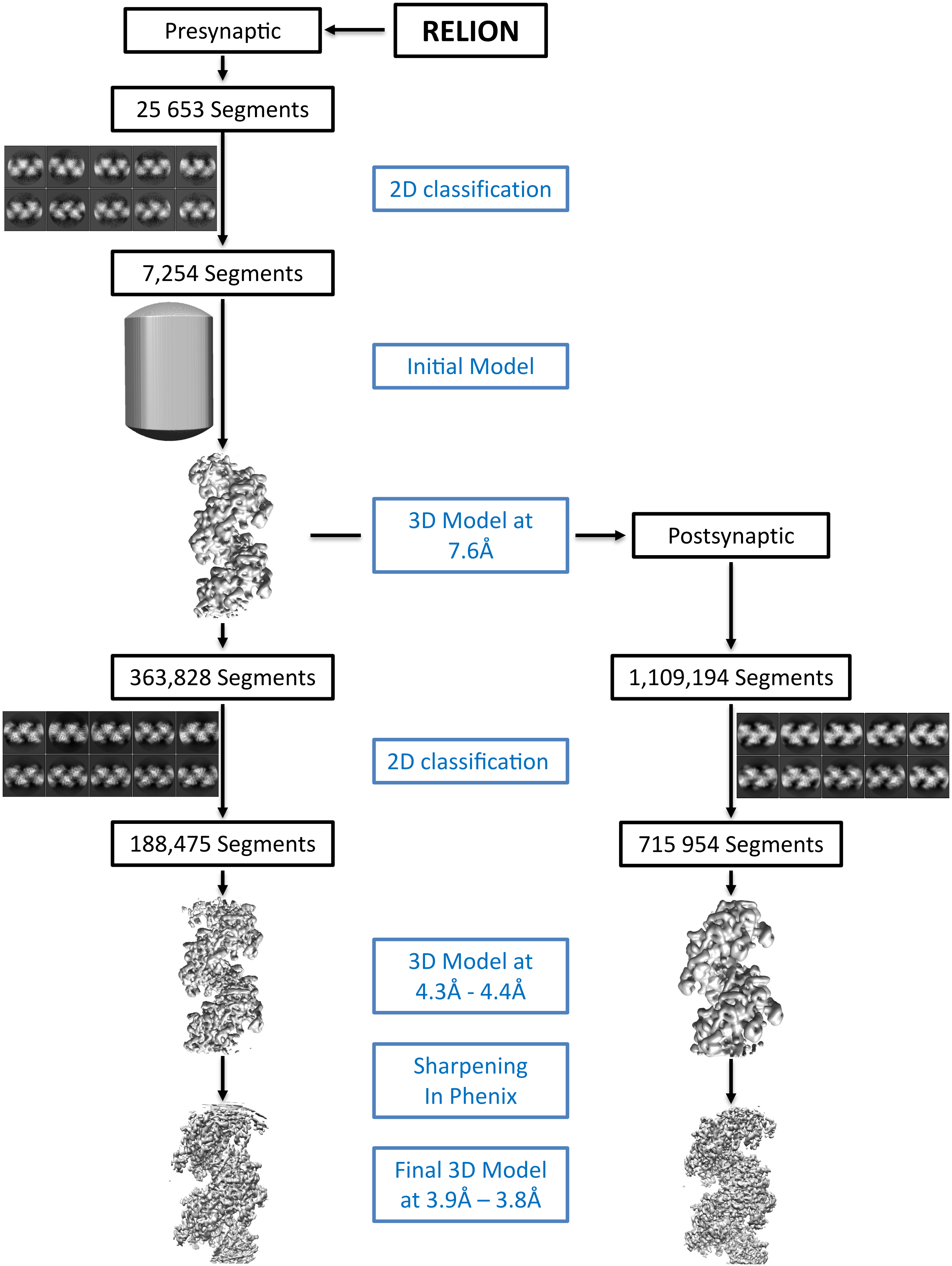
RELION
Presynaptic
25 653 Segments
2D classification
7,254 Segments
Initial Model
3D Model at
7.6Å
Postsynaptic
363,828 Segments
1,109,194 Segments
2D classification
188,475 Segments
715 954 Segments
3D Model at
4.3Å - 4.4Å
Sharpening
In Phenix
Final 3D Model
at 3.9Å – 3.8Å

**Supplemental Figure 3. Image processing in RELION 3.0.**

The flowchart summaries the image processing of filaments extracted from EM micrograph images. A starting reference model at 7.6Å was calculated by a quick processing of only 25 653 segments on the presynaptic complex, which permitted to determine the helical parameters and picked more effectively particles for the both complexes. Motion correction (MotionCor2) and local CTF correction (CTFFIND-4.1) software were also used during processing along in RELION. Several iterations of 3D refinement were carried out during the processing.

a.

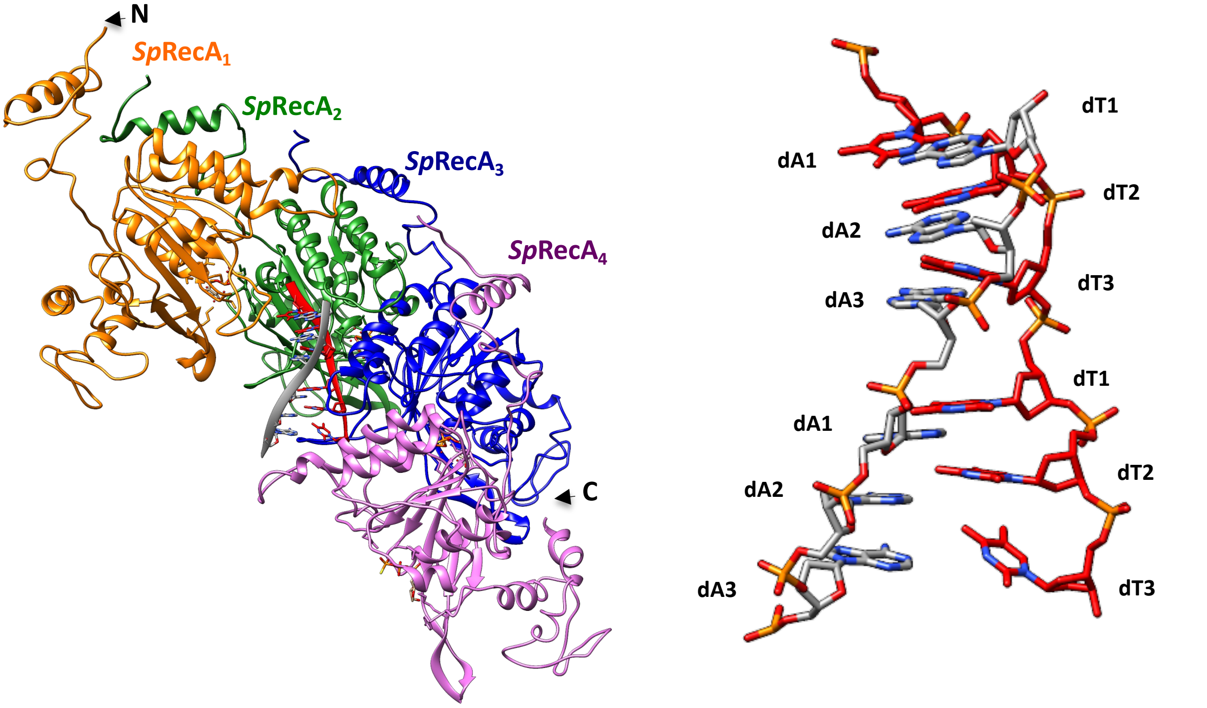
N
SpRecA1
SpRecA2
SpRecA3
SpRecA4
C
dT1
dA1
dT2
dA2
dT3
dA3
dT1
dA1
dT2
dA2
dA3
dT3

b.

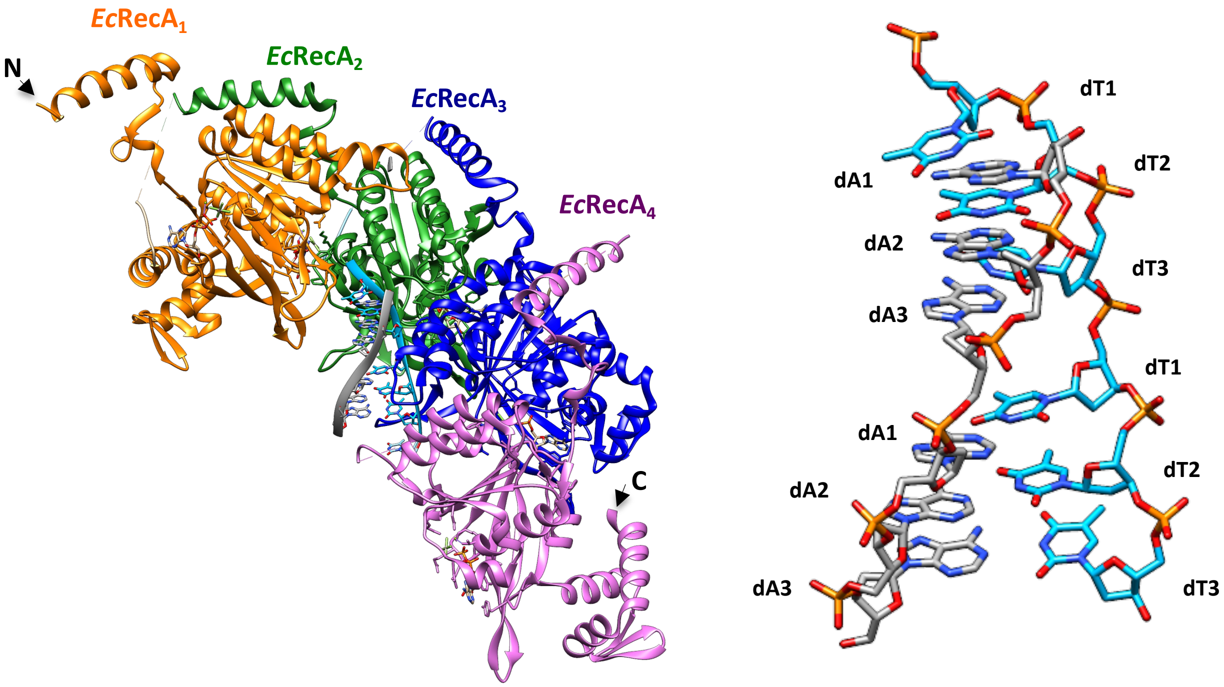
EcRecA1
N
EcRecA2
EcRecA3
EcRecA4
C
dT1
dT2
dA1
dA2
dT3
dA3
dT1
dA1
dT2
dA2
dA3
dT3

**Supplemental Figure 4. Structure comparison of the postsynaptic nucleoprotein filaments from *S. pneumoniae* and *E. coli.***

a and c. Structure of the RecA-ATPγS-dT-dA complex from *S. pneumoniae* (*Sp*RecA) (a) and *E. coli* (*Ec*RecA) (c). Four RecA protomers are numbered from the N-terminus of the first protomer to the C-terminus of the last protomer, coloured in orange, green, blue and purple respectively. A single stranded DNA (ssDNA) molecule composed of 8 thymidine nucleotides bound to *Sp*RecA and *Ec*RecA are represented in red and blue respectively. The complementary DNA strands are coloured in green. Four ATPγS molecules are shown in gold. b and d. Zoom on the double strand B-form DNA from *S. pneumoniae* (b) and *E. coli* (d) postsynaptic filaments. The dsDNA is numbered starting with the 5′-most nucleotide in each nucleotide triplet. The complementary DNA strand is juxtaposed to the primary strand in an antiparallel orientation. The two strands form a duplex with a complete set of Watson–Crick hydrogen bonds. The repeating unit is now a triplet of stacked base pairs, with adjacent base-pair triplets separated by a gap as in the presynaptic complex.

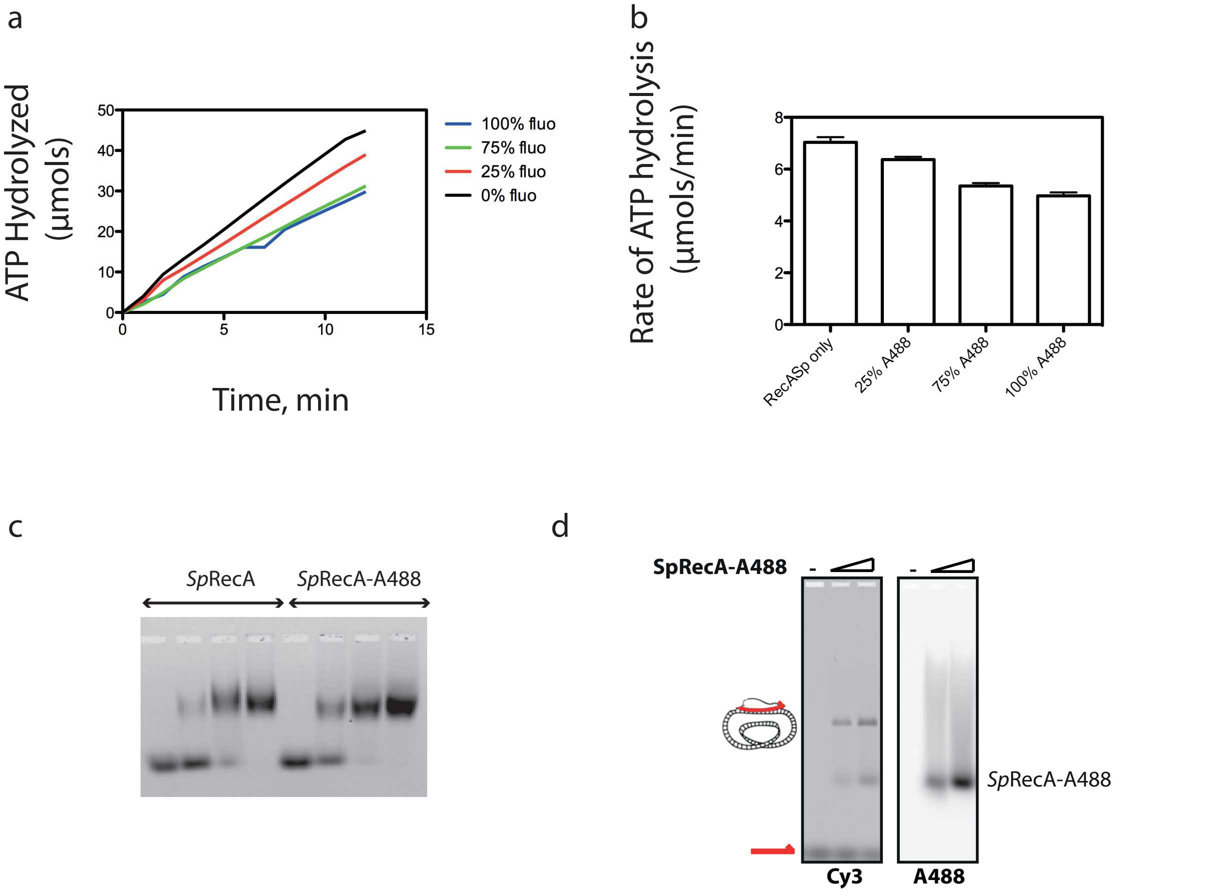


**Supplemental Figure 5**. a.b Comparisons of these ATP hydrolyzed rates for the different mix of proteins (*Sp*RecA/ *Sp*RecA$^{A488}$). The labeled protein hydrolyzed ATP upon ssDNA binding with a slight defect compared to the non-labeled protein. The experiment has been done in triplicate, 3 times. c. Native gel shift assay using agarose (1,2%, TB) gel and fluorescent labeled ssDNA probe (Size : 70 nucleotide length, probe : Cy3 ; concentration : 10 nM), in presence of non-hydrolysable analog of ATP (ATPγS); and the with the non-labeled *Sp*RecA protein (left) and the fluorescently-labeled *Sp*RecA$^{A488}$ protein (right) at increasing concentration: 0 , 250 nM, 500nM and 1µM. The ssDNA binding of the labeled and non-labeled proteins is similar in these experimental conditions. The experiment has been done 3 times. d. Deproteinized agarose gel of the D-loop reaction performed in presence cy3 oligonucleotide (100 mers) pUC18 vector, and of *Sp*RecA$^{A488}$.

a

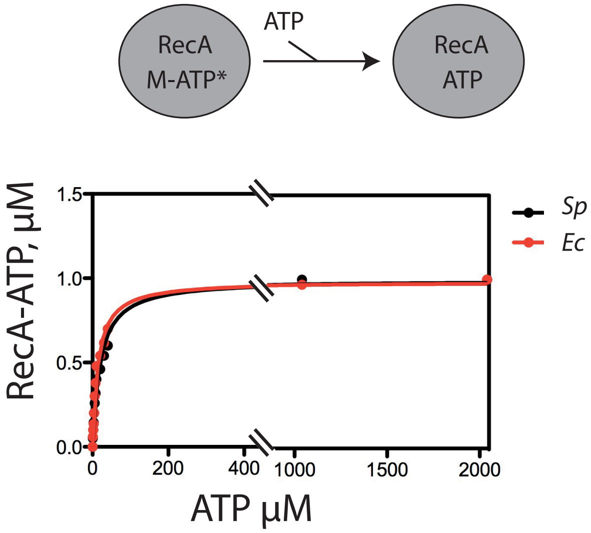


b

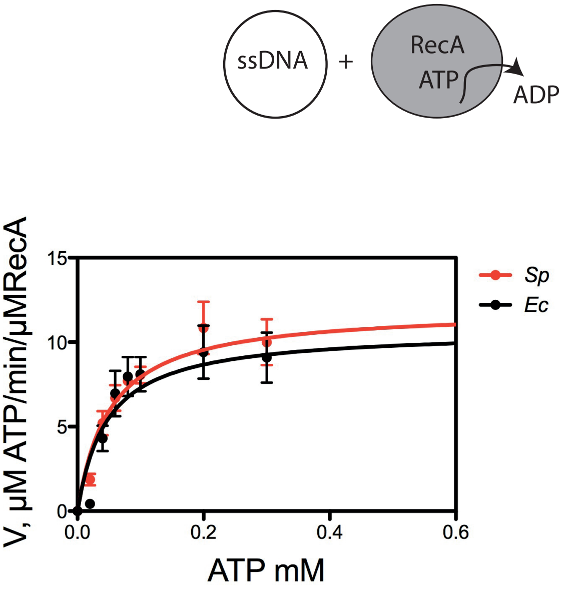


| Mg | *Sp* Km | *Sp* Vm | *Ec* Km | *Ec* Vm |
|---|---|---|---|---|
| 4mM | 0,05 +/-0,011 | 11,99 +/-0,8 | 0,047 +/-0,016 | 10,71 +/-1 |
| 1mM | 0,065 +/-0,007 | 21,74 +/-0,8 | 0,1254 +/-0,03 | 14,85 +/-1,4 |

**Supplemental Figure 6: ATP binding and ATP hydrolysis activities of *Sp*RecA and *Ec*RecA.**
a. *Sp*RecA and *Ec*RecA display similar ATP binding affinities (Kd of 19,98 µM +/-0,028 for *Ec*RecA and 14,58 µM +/- 0,023 for *Sp*RecA). The experiment was repeated in triplicate.
b. *Sp*RecA and *Ec*RecA display similar ATP hydrolysis activities.
Left: the data points represent the initial rates of ATP hydrolysis measured for *Sp*RecA (black) and *Ec*RecA (red) at the indicated concentrations of ATP The solid lines were generated using the Michaelis-Menten equation with Prism, GraphPad. The plots are the average of 3 experiments and the standard error of the mean value (+/- sem) is represented. Right: Parameters obtained from the fitted curves by the Michaelis Menten equation with Km in mM, and Vm in µM ATP/min/µM RecA.

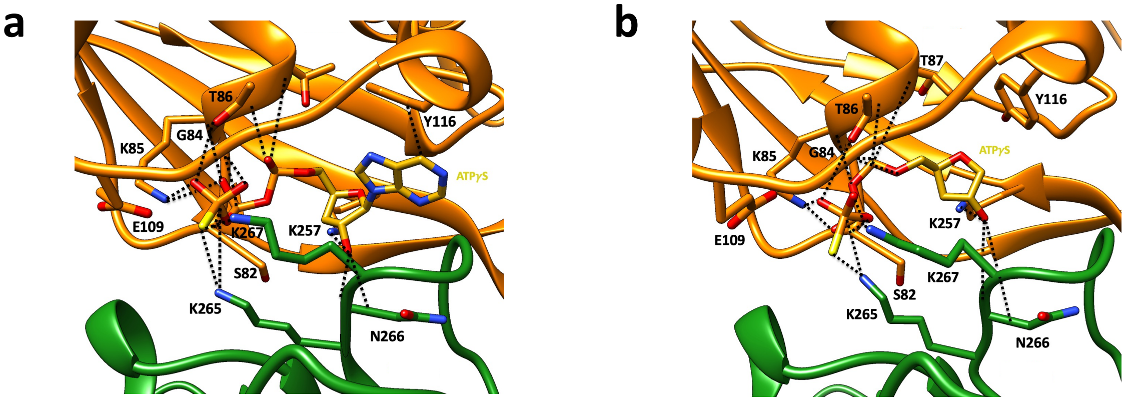


**Supplemental figure 7. RecA-ATP interface in the pre- and postsynaptic filaments from *S. pneumoniae*.**

a. Zoom of the RecA-ATPγS contacts in the presynaptic filament. The non-hydrolysable ATP analogue ATPγS binds at the interface of two protomers. The two RecA protomers are numbered and coloured as the Figure 2.a. Hydrogen-bond and covalent-bond interactions are shown by black dotted lines. b. Zoom of the RecA-ATPγS contacts at the interface of two protomers in the postsynaptic filament. The two RecA protomers are numbered and coloured as the Supplementary Figure 1.a. Hydrogen-bond and covalent-bond interactions are shown by black dotted lines.

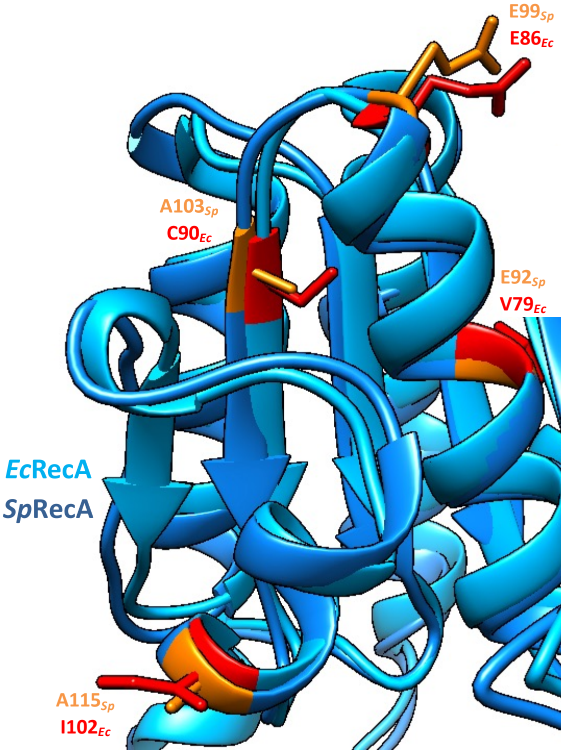


**Supplemental figure 8: RecA-SSB interface comparison in the pre- and postsynaptic filaments from *S. pneumoniae* and *E. coli*.**

Prominent residues in which amino acid changes bring about enhanced recombination potential in *E. coli* (from MM. Cox and coworkers) are shown in red, corresponding to amino acid A92, E99, A103 and A115 in *S. pneumoniae*, shown in orange. The *Sp*RecA and *Ec*RecA protomer are coloured respectively in deep blue and light blue.
